# Unsupervised changes in core object recognition behavior are predicted by neural plasticity in inferior temporal cortex

**DOI:** 10.1101/2020.01.13.900837

**Authors:** Xiaoxuan Jia, Ha Hong, James J. DiCarlo

## Abstract

Temporal continuity of object identity is a feature of natural visual input, and is potentially exploited -- in an unsupervised manner -- by the ventral visual stream to build the neural representation in inferior temporal (IT) cortex and IT-dependent core object recognition behavior. Here we investigated whether plasticity of individual IT neurons underlies human behavioral changes induced with unsupervised visual experience by building a single-neuron plasticity model combined with a previously established IT population-to-recognition-behavior linking model to predict human learning effects. We found that our model quite accurately predicted the mean direction, magnitude and time course of human performance changes. We also found a previously unreported dependency of the observed human performance change on the initial task difficulty. This result adds support to the hypothesis that tolerant core object recognition in human and non-human primates is instructed -- at least in part -- by naturally occurring unsupervised temporal contiguity experience.

## Introduction

Among visual areas, the inferior temporal (IT) cortex is thought to most directly underlie core visual object recognition in human and non-human primates (Ito, Tamura, Fujita, & Tanaka, 1995; Rajalingham & DiCarlo, 2019). For example, simple weighted sums of IT neuronal population activity can accurately explain and predict human and monkey core object recognition (COR) performance over dozens of such tasks (Majaj, Hong, Solomon, & DiCarlo, 2015). Moreover, direct suppression of IT activity disrupts COR behavior (Afraz, Boyden, & DiCarlo, 2015; Rajalingham & DiCarlo, 2019). These results were found in the face of significant variation in object latent variables including size, position and pose, and the high performance of the simple IT read-out (weighted sum) rests on the fact that many individual IT neurons show high tolerance to those variables (DiCarlo, Zoccolan, & Rust, 2012; Hung, Kreiman, Poggio, & DiCarlo, 2005; Li, Cox, Zoccolan, & DiCarlo, 2009), reviewed by (DiCarlo et al., 2012).

But how does the ventral stream wire itself up to construct these highly tolerant IT neurons? Simulated IT “neurons” in the deep layers of artificial neural networks (ANNs) have such tolerance, and provide quite accurate approximations of the adult ventral visual stream processing (Khaligh-Razavi & Kriegeskorte, 2014; Rajalingham et al., 2018; D. L. K. Yamins et al., 2014). However, those ANNs are produced by training with millions of supervised (labeled) training images, an experience regime that is almost surely not biologically plausible over evolution or post-natal development. That simple fact rejects all such ANNs as models of the construction of IT tolerance, regardless of whether or not the brain is executing some form of backpropagation-instructed plasticity (Plaut & Hinton, 1987). So the question remains open: how does the ventral stream wire itself up to *construct* a tolerant IT with minimal supervision?

The temporal stability of object identity under natural viewing (i.e. objects do not rapidly jump in and out of existence) has been proposed as a key available source of unsupervised information that might be leveraged by the visual system to construct neural tolerance, even during adulthood (Földiák, 1991; Hénaff, Goris, & Simoncelli, 2019; Rolls & Stringer, 2006; G. Wallis, Backus, Langer, Huebner, & Bulthoff, 2009; G Wallis & Bülthoff, 2001; Wiskott & Sejnowski, 2002). Consistent with this view, psychophysical results from human subjects show that unsupervised exposure to unnatural temporal contiguity experience (i.e. laboratory situations in which object *do* jump in and out of existence) reshapes position tolerance (Cox, Meier, Oertelt, & DiCarlo, 2005), pose tolerance (G Wallis & Bülthoff, 2001) and depth illumination tolerance (G. Wallis et al., 2009) as measured at the behavioral level.

Similarly, neurophysiological data from adult macaque IT show that unsupervised exposure to unnatural temporal contiguity experience reshapes IT neuronal position and size tolerance (Li & DiCarlo, 2008, 2010), in a manner that is qualitatively consistent with the human behavioral data.

Taken together, our *working hypothesis* is that the ventral visual stream is under continual reshaping pressure via unsupervised visual experience, that such experience is an important part of the construction of the tolerant representation that is ultimately exhibited at the top level of the ventral stream (IT), that the IT population feeds downstream causal mechanistic chains to drive core object discrimination behavior, and that the performance on each such behavioral tasks is well approximated by linear read-out of IT (Hung et al., 2005; Majaj et al., 2015).

However, there is a key untested element in this working hypothesis: is the single neuronal plasticity in adult monkey IT and the adult human behavioral changes resulting from unsupervised temporal contiguity experience quantitatively consistent with each other? In this study, we chose to focus on testing that missing link as it was far from obvious that it would hold up. In particular, the prior IT neurophysiology work was with basic level objects and produced seemingly large changes (~25% change in IT selectivity per hour of exposure in (Li & DiCarlo, 2010), and the prior human behavioral work was with subordinate level objects and produced significant, but subtle changes in behavior (e.g. ~3% performance change in (Cox et al., 2005). Moreover, if we found that the link did not hold, it would call into question all of the elements of the overall working hypothesis (esp. IT’s relationship to COR behavior, and/or the importance of unsupervised plasticity to the IT representation). Thus, either result would be important.

To test whether our *working hypothesis* is quantitatively accurate over the domain of unsupervised temporal contiguity induced plasticity, we sought to build a model to predict the changes in human object discrimination performance that should result from temporally-contiguity-experience-driven changes in IT neuronal responses. This model has three components: 1. a generative IT model (constrained by prior IT population response (Majaj et al., 2015)) that approximates the IT population representation space and can thus simulate the IT population response to any image of the objects (within the space) with variation in size; 2. an unsupervised plasticity rule (constrained by prior IT neural plasticity data (Li & DiCarlo, 2010)) to quantitatively describe and predict firing rate change of single IT neurons resulting from temporally-contiguous pair of experienced images and can thus be used to update the simulated IT population representation; 3. an IT-to-COR-behavior linking model (learned weighted sums, previously established by (Majaj et al., 2015)) to predict behavioral discrimination performance from the state of the IT (simulated) population both before and after each epoch of unsupervised experience.

To overcome the limitation of non-overlapping tasks in previous psychophysics and neurophysiology studies and to extend prior psychophysical work, we carried out new human behavioral experiments. Specifically, we measured the progression of changes in size-specific human object discrimination performance that resulted from unsupervised temporal contiguity experience using the same experience paradigm as the prior monkey neurophysiology work (Li & DiCarlo, 2010). We made these measurements for a wide range of object discrimination tasks, ranging from subordinate to basic level.

We found that – without any parameter tuning -- our overall model was quite accurate in its predictions of the direction, magnitude, and time course of the changes in human performance for all of the tested unsupervised experience manipulations. We also found that there was a strong dependency of learning effect on the initial task difficulty, with initially hard (d’<0.5) and initially easy (d’>2.5) COR tasks showing smaller measured learning effects than COR tasks of intermediate initial difficulty. The former was naturally explained by the overall model, and the latter was explained if we assumed a behavioral lapse rate of ~9% (Prins, 2012).

Taken together, this result shows that at least three separate types of studies (human unsupervised learning, IT unsupervised plasticity, and IT-to-COR-behavior testing) are all quantitatively consistent with each other. As such, this result adds support to the overall working hypothesis: that tolerant core object recognition is instructed -- at least in part -- by naturally occurring unsupervised temporal contiguity experience that gradually reshapes the non-linear image processing of the ventral visual stream without the need for millions of explicit supervisory labels (Krizhevsky, Sutskever, & Hinton, 2012; Y. LeCun et al., 1989; Riesenhuber & Poggio, 1999) and reviewed by (Yann LeCun, Bengio, & Hinton, 2015).

## Results

Our working hypothesis (see Introduction) predicts that IT population plasticity resulting from unsupervised visual experience should accurately predict the direction, magnitude, and time course of all changes in human object discrimination performance resulting from the same visual exposure. To quantitatively test these predictions, we first carried out a set of human psychophysical experiments with unsupervised experience that closely approximate an unsupervised experience paradigm that has been shown to reliably produce IT plasticity (measured as changes in size tolerance at single IT recording site) (Li & DiCarlo, 2010).

### Measure changes in human object discrimination performance induced by unsupervised visual experience

The basic experimental strategy is that, after testing initial object discrimination performance on a set of discrimination tasks (“*Test phase*”, Fig. 1a), we provide an epoch of unsupervised visual experience (“*Exposure phase*”, Fig. 1a) that is expected to result in IT plasticity (based on the results of (Li & DiCarlo, 2010)). At the end of the exposure epoch, we re-measure discrimination performance (*Test phase*), then provide the next epoch of unsupervised experience (*Exposure phase*), etc. (see Fig. 1a). This strategy allowed us to evaluate the accumulation of positive or negative behavioral changes (a.k.a. “learning”) resulting from four unsupervised experience epochs (400 exposure “trials” each) over approximately 1.5 to 2 hours. We include control discrimination tasks to subtract out any general learning effects.

**Figure 1.**
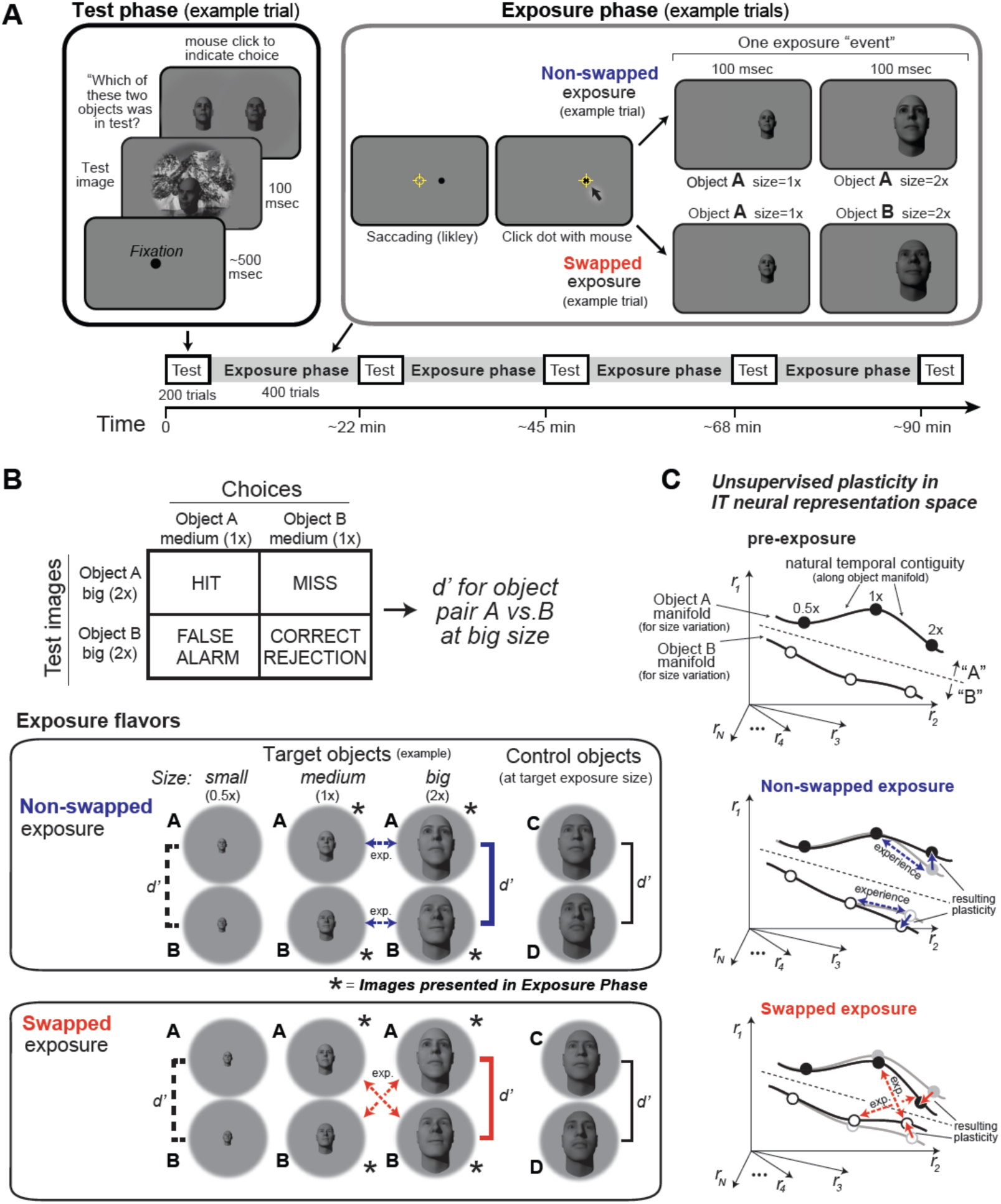
Experimental design and conceptual hypothesis. **A)** Illustration of human behavioral experimental design and an example trial from the Test Phase and from the Exposure Phase. **B)** Top: example confusion matrix for a 2AFC size-specific sub-task run during each Test Phase to monitor object-specific, size-specific changes in discrimination performance (see Methods). Bottom: the two unsupervised exposure flavors deployed in this study (see Methods). Only one of these was deployed during each Exposure Phase (see Fig. 2). Exposed images of example exposed objects (here, faces) are labeled with asterisks, and the arrows indicate the exposure events (each is a sequential pair of images). Note that other object and sizes are tested during the Test Phases, but not exposed during the Exposure Phase (see d’ brackets vs. asterisks). Each bracket with a d’ symbol indicates a pre-planned discrimination sub-task that was embedded in the Test Phase and contributed to the results (Fig. 2). In particular, performance for target-objects at non-exposed size (d’ labeled with dashed lines), target-objects at exposed size (d’ labeled with bold solid lines) and control objects (d’ labeled with black line) were calculated based on test phase choices. **C)** Expected qualitative changes in the IT neural population representations of the two objects that results from each flavor of exposure (based on Li et al., 2010). In each panel, the six dots show three standard sizes of two objects along the size-variation manifold of each object. Assuming simple readout of IT to support object discrimination (e.g. linear discriminant, see Majaj et al., 2015), non-swapped exposure tends to build size tolerant behavior by straightening out the underlying IT object manifolds, while swapped exposure tends to disrupt (“break”) size tolerant behavior by bending the IT object manifolds toward each other at the swapped size. This study asks if that idea is quantitatively consistent across neural and behavioral data without any parameter tuning.

**Figure 2.**
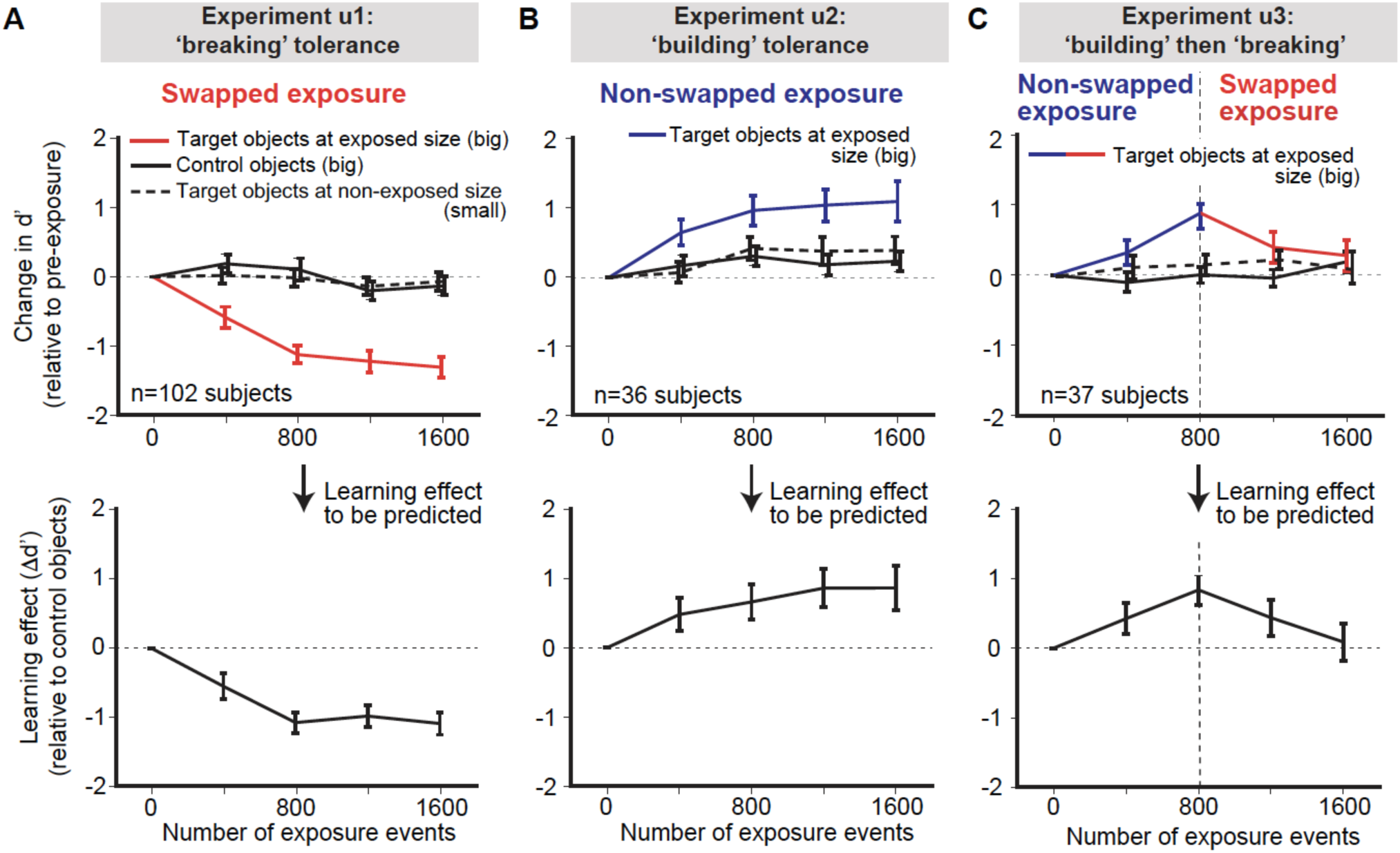
Measured human unsupervised learning effects as a function of amount of unsupervised exposure (each “exposure event” is the presentation of two, temporally-adjacent images, see Fig. 1A, right). We conducted three longitudinal unsupervised exposure experiments (referred to as u1, u2, u3). **A**) Swapped exposure experiment intended to “break” size tolerance (n=102 subjects; u1). Upper panels are the changes in d’ relative to initial d’ for targeted objects (faces) at exposed size (big) (red line), control objects (other faces) at the same size (big) (black line) and targeted faces at non-exposed size (small) (dashed black line) as a function of number of exposure events prior to testing. Lower panel is the to-be-predicted learning effect determined by subtracting change of d’ for control objects from the change of d’ for target objects (i.e red line minus black line). **B**) Same as A, but for non-swapped exposure experiment (n=36 subjects; u2). **C**) Same as A, except for non-swapped exposure followed by swapped exposure (n=37 subjects; u3) to test the reversibility of the learning. In all panels, error bars indicate bootstrapped standard error of the mean.

Specifically, we evaluated changes in discrimination performance (relative to initial performance) of each of a set of size-specific object discrimination tasks. A total of 174 human subjects on Amazon Mechanical Turk (see Methods and (Kar, Kubilius, Schmidt, Issa, & DiCarlo, 2019; Majaj et al., 2015; Rajalingham et al., 2018)) participated in this experiment.

To measure object discrimination performance in each subject, we used a set of 2-way alternative forced choice (2AFC) sub-tasks (size-specific object discrimination tasks; see Methods). These sub-tasks were randomly-interleaved (trial by trial) in each Test Phase, and the key test conditions used in the analyses (brackets indicated with d’s in Fig. 1B) were embedded within a balanced set of six sub-tasks and cover trials (see Fig. 1B and Methods).

Our first experiments used pairs of faces as the objects to discriminate, and we targeted our exposure manipulations at the big size (2x the baseline size; see Methods and Fig. 1; later, we targeted other pairs of objects and other sizes). Specifically, we used eight face objects from a previous study (Majaj et al., 2015). We chose these face objects at this size because, prior to unsupervised exposure, they had intermediate discriminability (mean d’ = 2.0±0.1 for big size, frontal view, n=28 pairs of faces), thus allowing us the possibility to measure both positive and negative changes in discrimination performance. For each subject, two target faces (manipulated during exposure) and two control faces (not shown during exposure) were randomly chosen from these eight faces.

Subjects were instructed to identify the single foreground face in a briefly presented test image (100ms) by choosing among two alternative choice faces immediately presented after the test image, one of which always correct (i.e. 50% chance rate). The test image contained one foreground object with variation in view (position, size, pose), overlaid on a random background (see Methods for test image generation). The choice images were always baseline views (i.e. size of ~2 degree, canonical pose) without background.

Similar to prior work testing the effects of unsupervised exposure on single-site IT recordings (Li & DiCarlo, 2010), each experiment consisted of two phases (Figure 1A): Test Phases to intermittently measure the size-specific object discrimination performance (d’) for the target face pair and control face pair (three d’ measured in each group of subjects, see Fig. 1B bottom); and Exposure Phases to provide unsupervised visual experience (pairs of images with different sizes in close temporal proximity; Figure 1A) that -- based on prior work -- was expected to improve or decrease the discrimination performance on the exposed objects.

The purpose of the Exposure Phase was to deploy unsupervised visual experience manipulations to target a particular object pair (two “target” objects) at particular views (e.g. sizes) of those target objects. For each exposure event, two images, each containing a different size object (frontal; no background) were presented consecutively (100 ms each) (see Methods for details). In non-swapped exposure events, both images contained the same object (expected to “build” size tolerance under the temporal contiguity hypothesis). In swapped exposure events, each images contained a different target object (expected to ‘break’ size tolerance under the temporal contiguity hypothesis). The conceptual predictions of the underlying IT neural population target object manifolds (DiCarlo & Cox, 2007) are that: non-swapped exposure events will straighten the manifold of each target object by associating size exemplars of the same object (as in the natural world), and that swapped exposure events will bend and decrease the separation between the two manifolds by incorrectly associating size exemplars of different objects (Fig 1C). This logic and experimental setup are adopted entirely from prior work (Li & DiCarlo, 2008, 2010).

In our studies here, we specifically focused on manipulating the size tolerance in the medium size (x1 of baseline view; ~2 deg) to big size (x2 of baseline view; ~4 deg) regime. Thus, the images shown during the Exposure Phase (indicated by * in Figure 1B) were always medium- and big-size, frontal-view of the target objects. We conducted three types of unsupervised exposure experiments (u): swapped (u1), non-swapped (u2) and non-swapped, followed by swapped (u3).

In Experiment u1 (swapped exposure events), we found that discrimination of the target face pair viewed at big size decreased with increasing numbers of exposure events (Figure 2A; top rows; red solid line; n=102 subjects). We found little to no change in performance for the non-exposed (small size) versions of those same faces (black dashed line; mean initial d’ is 1.2±0.1) or for non-exposed control faces (also tested at big size, black solid line). Lower panels in Figure 2A showed the learning effect defined by subtracting changes in control face discrimination performance (to remove general learning effects over the experience epochs, which turned out to be small; see Fig. 2A upper panel). In sum, we demonstrated an unsupervised, object-selective, size-selective temporal contiguity induced learning effect that was qualitatively consistent with prior work in ‘breaking’ tolerance (Cox et al., 2005; G Wallis & Bülthoff, 2001), and we measured the accumulation of that learning over increasing amounts of unsupervised exposure.

In Experiment u2 (non-swapped exposure events), we found that discrimination of the target face pair viewed at big size *increased* with increasing numbers of exposure events (Figure 2B; top rows; blue solid line; n=36 subjects). As in Experiment u1, we found little to no change in performance for the non-exposed (small size) versions of those same faces or for non-exposed control faces (also tested at big size, black solid line). This shows that, as predicted by the temporal contiguity hypothesis, unsupervised experience can build size tolerance at the behavioral level.

Interestingly, after ~800 exposure events, the exposure induced learning effects appeared to plateau in both ‘breaking’ tolerance conditions (Experiment u1, Fig. 2A) and ‘building’ tolerance conditions (Experiment u2), Fig. 2B, suggesting a limit in the measurable behavioral effects (see Discussion).

To test whether this unsupervised learning effect is reversible, we measured human performance in a combined design (Experiment u3) by first providing exposure epochs that should ‘build’ tolerance, followed by exposure epochs that should ‘break’ tolerance (n=37 subjects). Consistent with the results of Experiments u1 and u2, we found that size tolerance first increased with non-swapped (‘build”) exposures and then decreased with swapped (“break”) exposures (Fig. 2C), and that the effect did not spill over to the control objects.

In sum, these results confirmed that the effect of unsupervised visual experience was specific (to manipulated object and sizes) and strong even in adults. Furthermore, the measured human learning effect trajectories with different unsupervised visual exposure conditions (u1, u2, u3) were taken as behavioral effects that must -- without any parameter tuning -- be quantitatively predicted by our working hypothesis (that links IT neural responses to COR behavior; see Introduction). We next describe how we built an overall computational model to formally instantiate that working hypothesis to make those predictions.

### A generative model to simulate the population distribution of IT responses

To generate predictions of human behavior performance, we need to measure or otherwise estimate individual IT neural responses to the same images used in the human psychophysical testing (above) for a sufficiently large set of IT neurons (a.k.a. IT population responses). Because each of the objects we used in human psychophysics had been previously tested in neural recording experiments from monkey IT, we did not collect new IT population responses (very time consuming), but we decided instead to make suitably accurate predictions of the population pattern of IT response for test images of those objects. To do this, we built a generative model of the IT population based on the previously recorded IT population response to those objects. The output of this model is the firing rate of a simulated IT population to one presentation of a newly rendered test image (generated from the 64 base objects used in the previous study). With this model, we could simulate IT population responses to any image rendered from the psychophysically tested objects (approximately) without recording more neurons in behaving animals.

This generative IT model captures the IT neuronal representation space with a Multi-Dimensional Gaussian (MDG) model, assuming the distribution of IT population responses is Gaussian-like for each object (see Method for Gaussian validation) (Figure 3A). Because the MDG preserves the covariance matrix of IT responses to 64 objects, any random draw from this MDG gives rise to an object response preference profile (one response level for each of 64 objects) of a simulated IT neural site. To simulate the effect of changes in object size, for each simulated site, we randomly chose a size tuning kernel from a batch of size tuning curves that we had obtained by fitting curves to real IT responses across changes in presented object size (n=168 recording sites; data from (Majaj et al., 2015)). Motivated by prior work (Li et al., 2009), we assumed separability of object representation and size tuning, and simulated the response to any of the 64 objects at any of the three sizes as the outer product of the object and size tuning curves (Figure 3A bottom). However, since most measured size-tuning curves are not perfectly separable (DiCarlo et al., 2012; Rust & DiCarlo, 2010) and because the tested conditions included arbitrary background for each condition, we introduced independent clutter variance caused by backgrounds on top of this for each size of an object (Figure 3A) by randomly drawing from the distribution of variance across different image exemplars for each object. We then introduced trial-wise variance for each image based on the distribution of trial-wise variance of the recorded IT neural population. In sum, this model can generate a new, statistically typical pattern of IT response over a population of any desired number of simulated IT neural sites to different image exemplars within the representation space of 64 base objects at a range of sizes (here targeting “small”, “medium”, and “big” sizes to be consistent with human behavioral tasks; see Methods for details).

**Figure 3.**
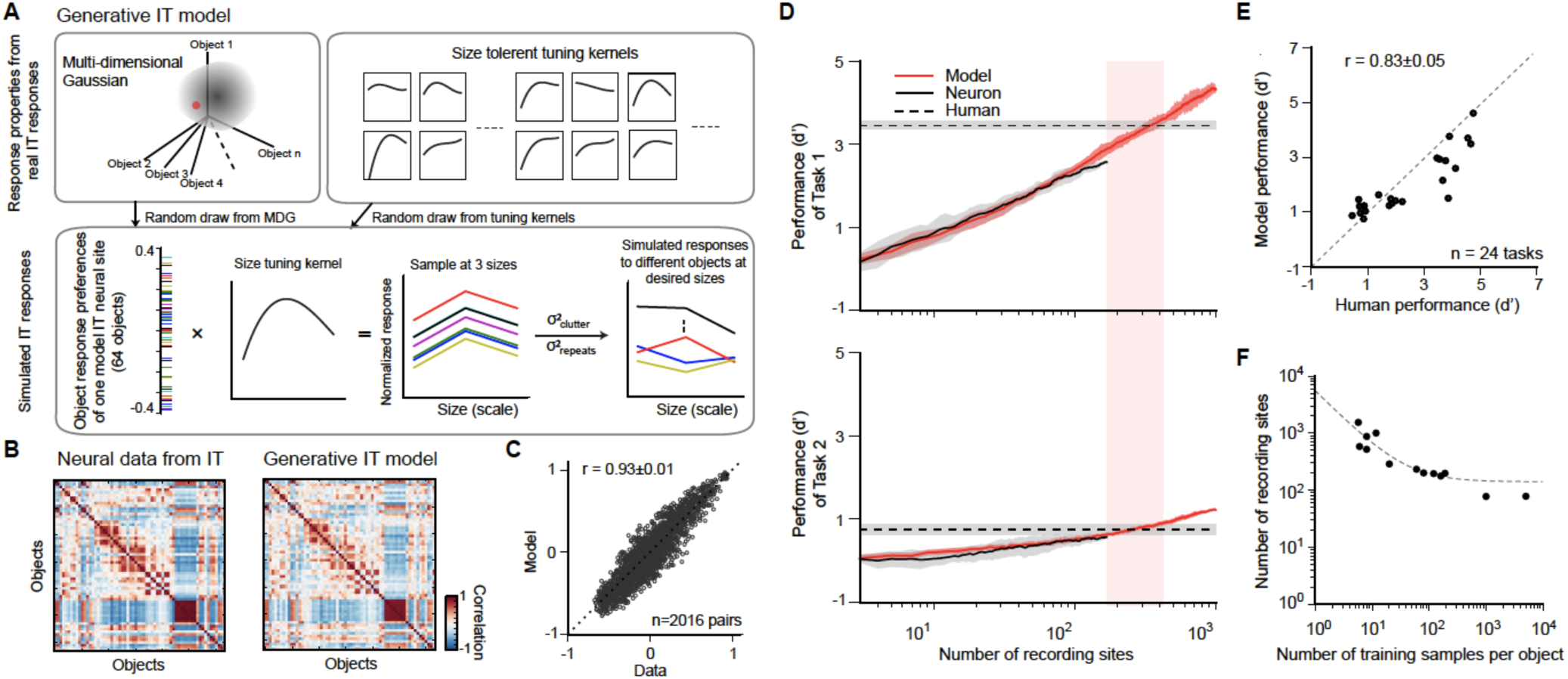
Generative IT model and validation of the IT-to-COR-behavior linking model. **A**) Generative IT model based on real IT population responses. Top left box: schematic illustration of the neuronal representation space of IT population with a multi-dimensional Gaussian (MDG) model. Each point of the Gaussian cloud is one IT neural site. Bottom box: The distribution of object preference of a simulated IT neural site for all 64 objects (i.e. a sample drawn from the MDG, highlighted as a red dot; each color indicates a different object). For each simulated site, a size tuning kernel is randomly drawn from the a batch of size tuning curves (upper right box) and multiplied by the object response distribution (outer product), resulting in a fully size-tolerant (i.e. separable) neural response matrix (64 objects x 3 sizes). To simulate the final mean response to individual images with different backgrounds, we added a “clutter” term to each element of the response matrix (σ^2^_clutter_; see Methods). To simulate the trial-by-trial “noise” in the response trials, we added a repetition variance (σ^2^_repeats_; see Methods). **B**) Response distance matrices for neuronal responses from real IT neuronal activity (n=168 sites) and one simulated IT population (n=168 model sites) generated from the model. Each matrix element is the distance of the population response between pairs of objects as measured by Pearson correlation (64 objects, 2016 pairs). **C**) Similarity of the model IT response distance matrix to the actual IT response distance matrix. Each dot represents the unique values of the two matrices (n=2016 object pairs), calculated for the real IT population sample and the model IT population sample (r=0.93±0.01). **D**) Determination of the two hyperparameters of the IT- to-behavior linking model. Each panel shows performance (d’) as a function of number of recording sites (training images fixed at m=20) for model (red) and real IT responses (black) for two object discrimination tasks (Task 1 is easy, human pre-exposure d’ is ~3.5; Task 2 is hard, human pre-exposure d’ is ~0.8; indicated by dashed lines). In both tasks, the number of IT neural sites for the IT-to-behavior decoder to match human performance is very similar (n~260 sites), and this was also true for all 24 tasks (see E), demonstrating that a single set of hyperparameters (m=20, n=260) could explain human pre-exposed performance over all 24 tasks (as previously reported by Majaj et al., 2015). **E**) Consistency between human performance and model IT based performance of 24 different tasks for a given pair parameters (number of training samples m = 20 and number of recording sites n = 260). The consistency between model prediction and human performance is 0.83±0.05 (Pearson correlation +/- SEM). **F**) Manifold of the two hyperparameters (number of recording sites and number of training images) where each such pairs (each dot on the plot) yields IT-based performance that matches initial (i.e. pre-exposure) human performance (i.e. each pair yields a high consistency match between IT model readout and human behavior, as in E). The dashed line is an exponential fit to those dots.

To check if the simulation is statistically accurate in terms of the layout of images in IT population representation space, we compared the representation similarity matrix (RSM; correlation between neuronal population responses to different images) of different draws of a simulated IT with the RSM measured from the actual IT neural data (Figure 3B). One typical example of that is shown in Figure 3C, revealing high correlation of the two RSMs (r=0.93±0.01). While this does not guarantee that any such simulated IT population is fully identical to an IT population that might exist in an actual monkey or human IT, our goal was simply to get the simulated IT population response distribution in the proper range (up to second order statistics).

### A standard IT-to-COR-behavior linking model for core object discrimination behavior

To make predictions about how IT neural changes will results in behavioral changes, we first needed a model to establish the linkage between IT population response and core object discrimination behavior prior to any experience-induced effects. We have previously found that simple weighted linear sums of IT neural responses accurately predict the performance (d’) of human object discrimination for new images of those same objects (here termed the IT-to-COR-behavior linking model) (Majaj et al., 2015). That model has only two free hyperparameters: the number of neural sites and the number of labeled (a.k.a. “training”) images used to set the weights of the decoder for each object discrimination. Once those two hyperparameters are locked, it has been empirically demonstrated that performance for any object discrimination task on new images is accurately predicted by its trained decoder (Majaj et al., 2015). To test whether the simulated IT population activity from the generative IT model (above) could quantitatively reproduce those prior results and to lock these two hyperparameters, we compared the predicted performance (for any given object recognition task) based on the simulated IT population (Figure 3D; red solid line) with the predicted performance based on the previously recorded IT neural population (black solid line). We did this as a function of number of recording sites, for a set of object recognition tasks. Figure 3D illustrated two example tasks (errorbar is standard error across 40 random subsamples of recording sites). As expected, we found that the model predictions overlapped with decoded performance of real IT neural sites, indicating that our generative IT model has captured the relevant components of the IT population response.

We next set out to choose the two free hyperparameters (number of sites and number of training examples). The crossing point with human performance in Figure 3D reflects how many neural sites are necessary to reach human performance level for a given number of training samples. Unlike the real IT neural data (n=168 recording sites) which required extrapolation to estimate the number of sites matching human absolute performance (Majaj et al., 2015), we simulated up to 1000 IT sites with the generative model to cover the range of neural sites necessary to reach human performance.

Consistent with Majaj et al (Majaj et al., 2015), we found that the number of simulated IT sites required to match human was similar across different tasks (260±23 IT sites given 20 training images (tested over 24 object discrimination tasks: low variation 8-way tests: 8 basic level, 8 car identification and 8 face identification tasks; previously used in (Majaj et al., 2015)). Specifically, we here used 260 sites with 20 training samples for all tasks, and the match between the decoded simulated IT performance and human performance over all discrimination tasks was r=0.83±0.05 (n=24 tasks), similar to previously reported match between decoded neural IT performance and human for the same tasks (r=0.868 from (Majaj et al., 2015)). Note that other specific combinations of the number of IT sites and the number of training examples are also suitable (Figure 3F) and we explored this later.

In sum, by setting the two decoder hyperparameters to match initial human performance, we established a fixed linear decoder rule that could be applied to our simulated IT population to quantitatively predict the expected performance of the subject (i.e. the owner of that IT population) for any object discrimination task.

The consequence is that, because the linkage model between the IT population and behavior is now fixed, any changes in the IT population are automatically mapped to predicted changes (if any) in behavioral performance (with no free parameters). From here on, we locked down the generative IT model and the decoders that matched human initial performance (before learning), and we combine both of these models later to make predictions of direction and magnitude of behavioral performance change (if any) that should result from any given change in the IT population (Fig. 2).

### Unsupervised IT plasticity rule

To model the IT neural population response changes that result from the unsupervised visual experience provided to the human subjects, we developed an unsupervised IT plasticity rule guided by previous studies of IT plasticity effects in the rhesus monkey that used the same paradigm of unsupervised visual experience that we provided here to our human subjects (Li et al., 2009; Li & DiCarlo, 2008, 2010). In particular, we set out to build an unsupervised IT plasticity rule that could predict the (mean) response change that occurs in each and every IT neuron as a result of each presented pair of temporally-contiguous visual images. We assumed that the same model would also apply to human “IT” without any parameter modifications (see Discussion).

Those prior monkey studies revealed that exposure to altered (“swapped”) visual statistics typically disrupts the size-tolerance of single IT neurons, while exposure to normal statistics in visual experience (non-swapped condition) typically builds size tolerance (Li & DiCarlo, 2010). To develop our unsupervised IT plasticity rule, we replicated the exact same experiment used in the monkeys on simulated-IT neural sites. Figure 4A illustrates the exposure design for single IT sites, where the preferred object (P) and non-preferred object (N) of each neural site are defined by the magnitude of neuronal activity (z-scored across all objects for each IT site). Selectivity of a neural site is measured by the difference of neuronal responses to its preferred and non-preferred objects (P-N) /(P+N), the same as (Li & DiCarlo, 2010).

**Figure 4.**
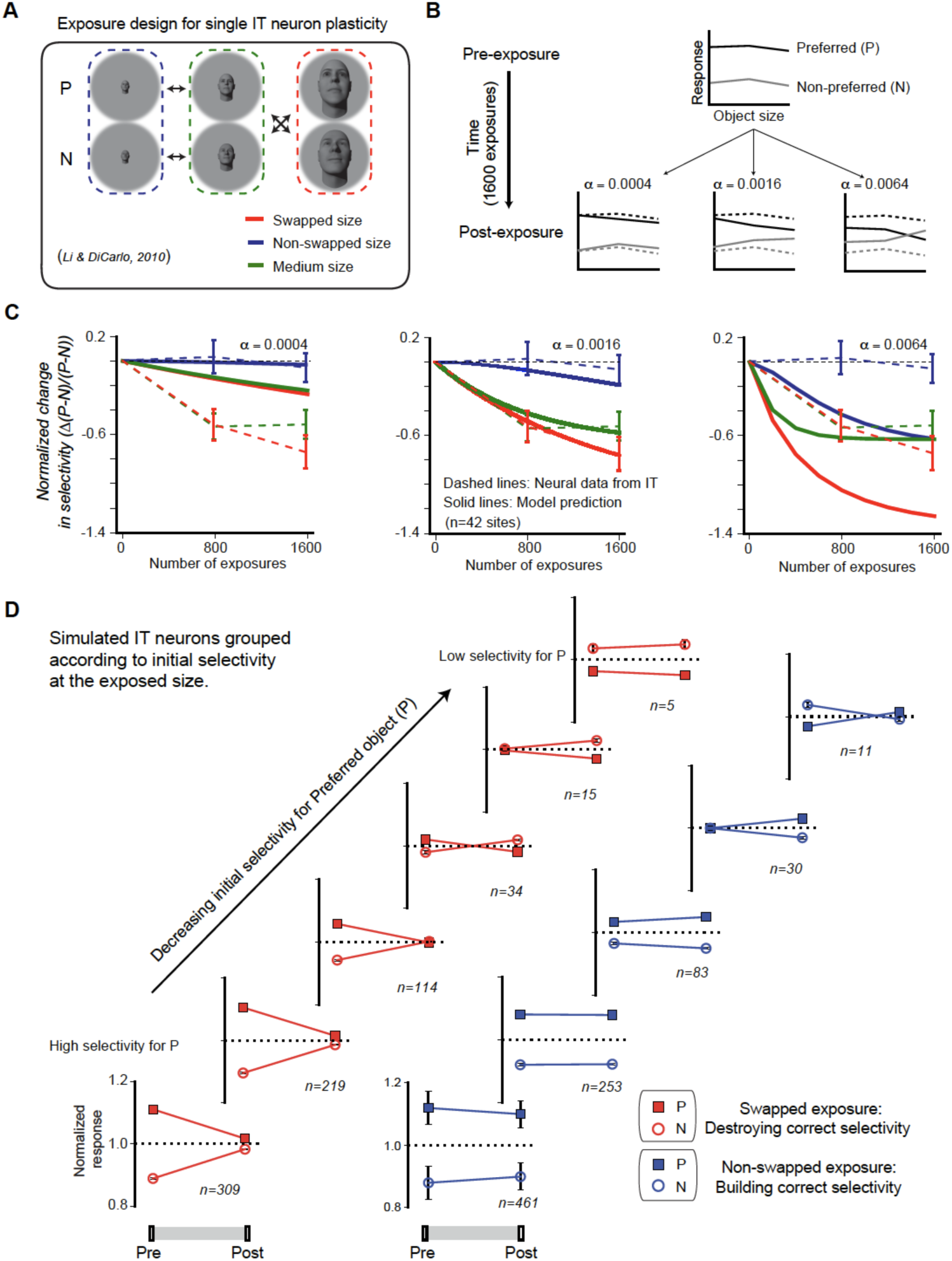
Temporal continuity-based IT neuronal plasticity rule. **A**) Illustration of exposure design for IT neuronal plasticity (adapted directly from (Li & DiCarlo, 2010)) measured with single electrodes. P refers to preferred object of the IT unit, N refers to non-preferred object of that unit. **B**) We developed an IT plasticity rule that modifies the model neuronal response to each image in individual neural sites according to the difference in neuronal response between lagging and leading images for each exposure event (see Methods). The figure shows the model predicted plasticity effects for a standard, size-tolerance IT neural site and 1600 exposure events (using the same exposure design as Li and DiCarlo, 2010; i.e. 400 exposure events delivered (interleaved) for each of the four black arrows in Panel A) for three different plasticity rates. Dashed lines indicate model selectivity pattern before learning for comparison. **C**) Normalized change over time for modeled IT selectivity for three different plasticity rates. Dashed lines are the mean neuronal plasticity results from the same neural sites in Li and DiCarlo, 2010 (mean change in P vs. N responses, where the mean is taken over all P > N selective units that were sampled and then tested, see (Li & DiCarlo, 2010)). Solid lines are the mean predicted neuronal plasticity for the mean IT model “neural sites” (where these sites were sampled and tested in a manner analogous to (Li & DiCarlo, 2010), see Methods). Blue line indicates the change in P vs. N selectivity at the non-swapped size, green indicates change in selectivity at the medium size and red indicates change in selectivity at the swapped size. Error bars indicate standard error of the mean. **D**) Mean swapped object (red) and non-swapped object (blue) plasticity that results for different model IT neuronal sub-groups – each selected according their initial pattern of P vs. N selectivity (analogous to the same neural sub-group selection done by (Li & DiCarlo, 2010) c.f. their Fig. 6 and 7).

We used a Hebbian-like (associative) plasticity rule (Caporale & Dan, 2008; D.O. Hebb, 1949; Oja, 1982; Paulsen & Sejnowski, 2000), which updates firing rate for each pair of images based on the difference of neuronal firing rate (FR) between the lagging and leading images (see Method). Our plasticity rule states that the modification of firing rate of each IT unit to the leading image at time t is equal to the difference of firing rate between lagging and leading images multiplied by a plasticity rate α. This plasticity rule tends to reduce the difference in neuronal responses to consecutive images and implies a temporal association to images presented close in time. The plasticity rule is conceptually similar to previously proposed temporal continuity plasticity or (a.k.a. slow feature analysis) (Berkes & Wiskott, 2005; Földiák, 1990, 1991; Mitchison, 1991; Sprekeler, Michaelis, & Wiskott, 2007). It is physiologically attractive because the findings on short-term synaptic plasticity revealed that synaptic efficacy changes over time in a way that reflects the history of presynaptic activity (Markram, Gerstner, & Sjöström, 2012; Markram, Lübke, Frotscher, & Sakmann, 1997). Even though conceptually similar, our plasticity rule is a ‘descriptive’ rather than a ‘mechanistic’ rule of plasticity at all stages of the ventral stream. That is, the rule does not imply that all the underlying plasticity is in IT cortex itself – but only aims to quantitatively capture and predict the changes in IT responses resulting from unsupervised visual experience. It is general in a sense that it can make predictions for different objects or dimensions of variations, but it is (currently) limited in that it: only applies to temporally paired image associations, it ignores any correlation in the neural response patterns, and it assumes that updates occur only in the responses to the exposed images (i.e. non-exposed object/size combinations are not affected).

To show the effects of this unsupervised IT plasticity rule, we illustrate with an example simulated IT neural site. The simulated neural site in Figure 4B was initialized to be – like many adult monkey IT neurons – highly size tolerant: its response to a preferred object (P) is always greater than response to a non-preferred object (N) at each size. After applying the unsupervised exposure design in Figure 4A (200 exposure events for each arrow, 1600 exposure events in total), the responses to each of the six conditions (2 objects x 3 sizes) evolved as shown in Figure 4B. We note two important consequences of this plasticity rule. First, because the rule was designed to decrease the difference in response across time, responses to images presented consecutively tend to become more similar to each other, which results in a reduction in the response difference between P and N at both the swapped and the non-swapped sizes. Second, once the neural site reached a state in which its response is no different over consecutively exposed images, the learning effect saturates. Notably, unlike adaptive changes in plasticity rate in the typical supervised optimization of deep neural networks (Kingma & Ba, 2014), our plasticity rate is kept constant over the “lifetime” of the model. The gradual shrinkage of learning effect (Δ(P-N) /(P+N)) as more and more exposure events are provided was a consequence of the gradual reduction in the absolute difference between neuronal responses to the two consecutive images that makeup each exposure event.

There is only one free parameter in our plasticity rule equation--the plasticity rate α. We determined this parameter using the single electrode physiology data collected previously in the lab (Li & DiCarlo, 2010). Figure 4C shows the average IT plasticity effect that results from different settings of α (here the plasticity effect is defined by the normalized changes in selectivity: Δ(P-N)/(P-N), exactly as was done in (Li & DiCarlo, 2010). As expected, a higher plasticity rate (α) results in greater model IT plasticity effects (Fig 4C). We chose the plasticity rate (α) that best matched the prior monkey IT neurophysiology results (i.e. the α that resulted in the minimal difference between the model IT plasticity effect (solid lines) and the experimentally reported IT plasticity effect (dashed lines) for swapped, non-swapped and medium object sizes; see Fig 4C middle). The best α is 0.0016 nru per exposure event (nru=normalized response units; see Methods for intuition about approximate spike rate changes). Once we set the plasticity rate, we locked it down for the rest of this study (otherwise noted later where we test rate impact).

We next asked if our IT plasticity rule naturally captured the other IT plasticity effects reported in the monkey studies (Li & DiCarlo, 2010). Specifically, it was reported that, for each neural site, the selectivity that results from a fixed amount of unsupervised exposure depends on the initial selectivity of that site. Thus, the unsupervised “swapped” experience manipulation causes a reduction of selectivity for neural sites that show a moderate level of initial P (preferred object) vs. N (non-preferred object) selectivity at the swapped size, and the same amount of unsupervised experience *reverses* the selectivity of neuronal sites that show a low level of initial selectivity at the swapped size (i.e. cause the site to, oxymoronically, prefer object N over object P). Li & DiCarlo (2010) also reported that the more natural, “non-swapped” experience manipulation caused a *building* of new selectivity (for neuronal units that initially show a strong preference for P at some sizes, but happened to have low P vs. N selectivity at the non-swapped size).

We tested for both of these effects in our model by selecting subsets of neural sites in the simulated IT population in exactly the same way as (Li & DiCarlo, 2010) (sampled from n=1000 simulated IT units) and then applied the plasticity rule to those units. We found a very similar dependency of the IT plasticity to those previously reported IT plasticity effects (Figure 4D; cf. Fig. see Figures 6 and 7 of (Li & DiCarlo, 2010)).

**Figure 5.**
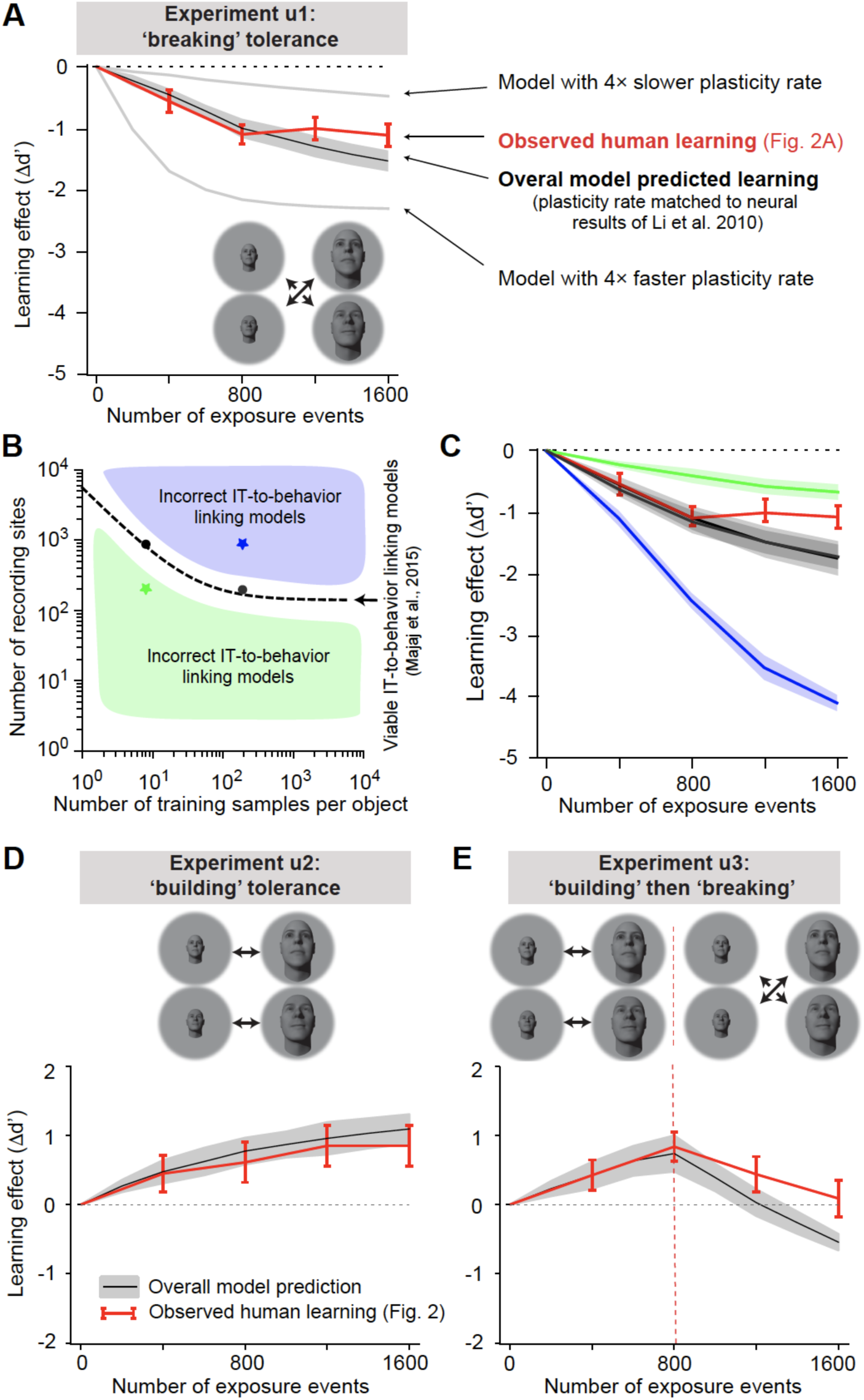
Overall model predicted learning effects vs. actual learning effects (red). **A**) Overall model predicted learning effect (solid black line) for experiment u1 (swapped exposure) with the IT-to-behavior linking model matched to initial human performance (hyperparameters: number of training images m = 20, number of model neural sites n = 260; see Fig. 3) and the IT plasticity rate matched to prior IT plasticity data (0.0016, see Fig. 4). Red line indicates measured human learning effect (reproduced from Fig. 2A lower). Grey lines indicate model predictions for 4 times smaller plasticity rate and 4 times larger plasticity rate. Error bars are standard error over 100 runs of the overall model, see text. **B**) Decoder hyperparameter space: number of training samples and number of neural features (recording sites). The dashed line indicates pairs of hyperparameters that give rise to IT IT-to-behavior performances that closely approximate human initial (pre-exposure) human object recognition performance over all tasks. **C**) Predicted unsupervised learning effects with different choices of hyperparameters (in all cases, the IT plasticity rate was 0.0016 – i.e. matched to the prior IT plasticity data, see Fig. 4). The two black lines (nearly identical, and thus appear as one line) are the overall model predicted learning that results from hyperparameters indicated by the black dots (i.e. two possible correct settings of the decoder portion of the overall model, as previously established by (Majaj et al., 2015)). Green and blue lines are the overall model predictions that result from hyperparameters that do not match human initial performance (i.e. non-viable IT-to-behavior linking models). **D**) Predicted learning effect (black line) and measured human learning effect (red) for building size tolerance exposure. **E)** Model predicted learning effect (black line) and measured human learning effect (red) for building and then breaking size tolerance exposure. In both D and E, the overall model used the same parameters as in A (i.e. IT plasticity rate of 0.0016, number of training samples m = 20 and number of model neural sites n = 260).

**Figure 6.**
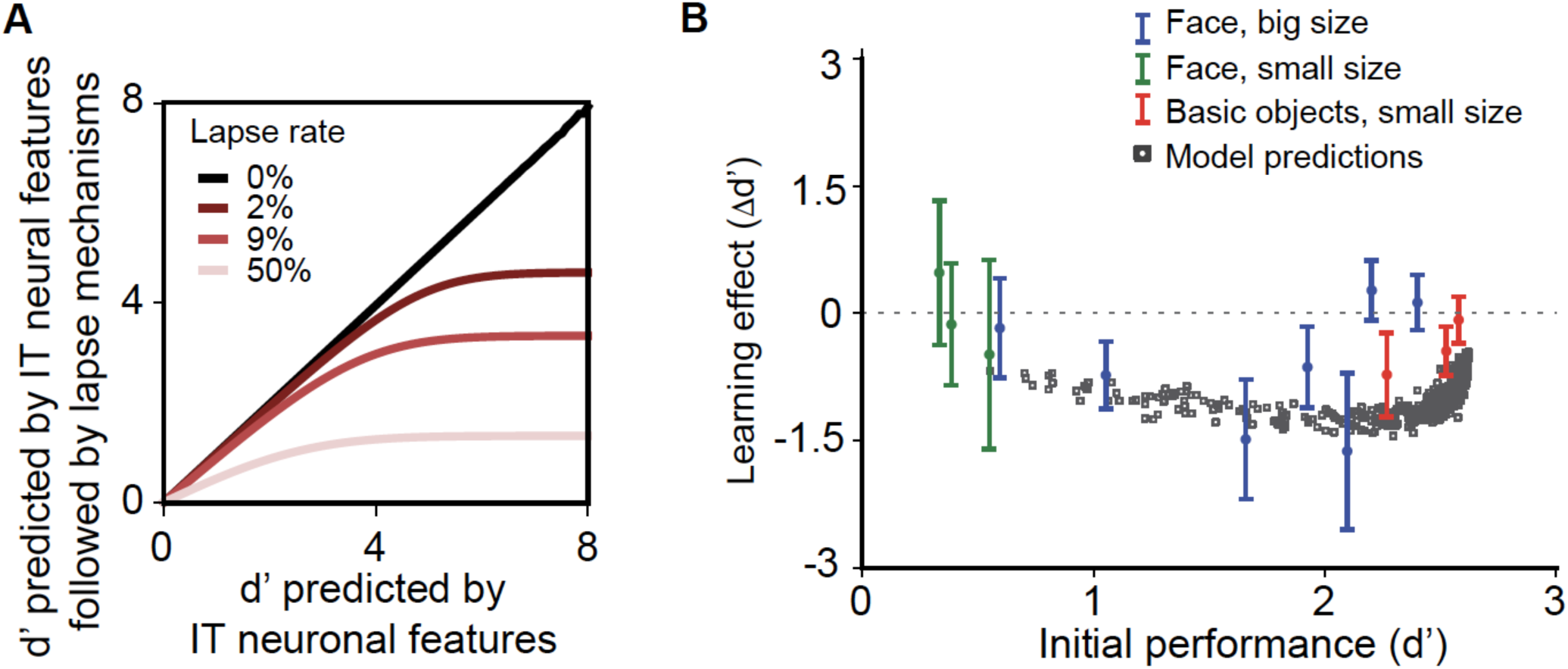
Learning effect as a function of initial task difficulty. A) Illustration of the saturation of measured d’ that results from the assumption that the subject guesses on a fraction of trials (lapse rate), regardless of the quality of the sensory evidence provided by the visually-evoked IT neural population response (x axis). B) Measured human learning effect for different tasks (colored points) as a function of initial (pre-exposure) task difficulty (d’) with comparison to model predictions (grey squares; model lapse rate for this plot is 9%). Different colors indicate different types of tasks and exposures: green indicates small-size face discrimination learning effect induced with medium-small swapped exposure (n=100 subjects); blue indicates big-size face discrimination learning effect induced with medium-big swapped exposure (n=161 subjects); red indicates small-size basic level discrimination learning effect induced with medium-small swapped exposure (n=70 subjects). Error bars indicate bootstrapped standard error of the mean over subjects (n~20-40 subjects per point).

Given that our IT plasticity rule tends to pull the response of temporally contiguous images toward each other (Berkes & Wiskott, 2005; Földiák, 1990, 1991; Mitchison, 1991; Sprekeler et al., 2007), it is not entirely obvious how this can build selectivity (i.e. pull response to P and N apart). The reason this occurs is that some IT neural sites have (by chance draw from the generative model of IT, above) initially high selectivity for P vs. N at the medium size and no selectivity at (e.g.) the big size. (Indeed, such variation in the IT population exists as reported in (Li & DiCarlo, 2010)). By design, the non-swapped (“natural”) unsupervised exposure temporally links P_med_ (high response) with P_big_, which – given the plasticity rule -- tends to pull the P_big_ response upward (pull it up higher than N_big_). In addition, the non-swapped exposure links N_med_ (low response) with N_big_, which can pull the N_big_ response downward (provided that the N_med_ response is initially lower than the N_big_ response). Both effects thus tend to increase the P_big_ vs. N_big_ response difference (that is, both effects tend to “build” selectivity for P vs. N at the big presentation size, which results in the neural site preferring object P over object N at both the medium and the big sizes – a property referred to as size “tolerance”). This effect is observed in single IT neural site size tuning curve for P and N before and after learning (see Figure 3 in (Li & DiCarlo, 2010)). Indeed, it is this effect that conceptually motivated temporal contiguity plasticity in the first place -- natural-occurring statistics can be used to equalize the responses to the same object over nuisance variables (such as size).

In sum, our very simple IT plasticity rule quantitatively captures the average IT plasticity effects for which its only free parameter was tuned, and it also naturally captures the more subtle IT neural changes that have been previously described.

### Putting together the overall model to predict human unsupervised learning effects

To summarize, we have: A. built and tested a generative IT model that captured the object representation space and variability in the actual primate IT population; B. locked down a set of parameters of a linear decoder rule that quantitatively links the current state of the simulated IT population to predicted human performance on any discrimination task (including the ones we plan to test); C. defined an IT plasticity rule that describes how each individual IT neural site changes as a result of each unsupervised exposure event and we locked down the only free parameter (plasticity rate) in that rule to match existing monkey IT plasticity data (see Supp. Fig. 1A). At this point, we could – without any parameter tuning -- combine each of these three model components into a single overall model that predicts the direction, magnitude and time course of human unsupervised learning effects that should result from any unsupervised learning experiment using this exposure paradigm (pairwise temporal image statistics).

Specifically, to generate the predictions for each of unsupervised learning experiments (u: u1, u2, u3, see Fig. 2) we: 1) initialized a potential adult human IT (from the generative IT model) with a total of 260 simulated IT recording sites, 2) built linear decoders for the planned object discrimination tasks that read from all 260 sites, using 20 training examples for each and every task, 3) froze the parameters of all such decoders (i.e. froze the resulting weighting on each simulated IT neural site on the “subject’s” object choice decision), 4) “exposed” the IT model population to the same unsupervised temporal exposure history as the human subjects, using the IT plasticity rule to update the model “IT” after each temporally-adjacent image exposure pair to update the responses of each simulated IT neural site (note that the plasticity trajectory of each neural site is dependent on both its initial object/size response matrix (1), and the sequence of images applied during unsupervised experience (u)), 5) measured the changes in “behavioral” performance of the overall model (changes in object discrimination performance of the (frozen) decoders (2)), and 6) took those changes as the predictions of the expected changes in human performance that should result from that unsupervised experience (u).

To give robust model estimates of the average predicted effects, we repeated this process (1-6) 100 times for each experiment (u) and averaged the results, which is analogous to running multiple subjects and averaging their results (as we did with the human data, see Fig. 2). For clarity, we note that the prediction stochasticity is due to: random sampling of the IT generative population, the clutter variability introduced in the generative IT model when generating the initial population response for each test image, the trial-by-trial variation in the simulated IT responses, the random unsupervised exposure event sequence (see Methods), and randomly drawn test images, all of which we expect to average out.

Note that, in expecting that these overall model predictions might be accurate, we are implicitly making the following assumptions: a) monkey IT and human IT are approximately identical (Kriegeskorte et al., 2008; Rajalingham, Schmidt, & DiCarlo, 2015), b) the linkage of IT to behavioral performance is approximately identical (as suggested by (Majaj et al., 2015); c) human IT unsupervised plasticity is the same as monkey IT unsupervised plasticity, and d) humans do not re-learn or otherwise alter the assumed mechanistic linkage between IT and behavior during or after unsupervised visual experience (at least not at the time scales of these experiments: 1.5 to 2 hours).

### Results: predicted learning vs. observed learning

Figure 5A,D,E show the model-predicted learning effects (black solid line) for each of the three unsupervised experiments (u1, u2, u3) plotted on top of the observed measured human learning effects (red line, reproduced from the learning effects shown in Fig. 2 bottom). For each experiment, we found that the overall model did a very good job of predicting the direction, magnitude and time course of the changes in human behavior. The directional predictions are not surprising given prior qualitative results, but the accurate predictions of the magnitude and time course are highly non-trivial (see below). Despite these prediction successes, we also noticed that the predictions were not perfect, most notably after large numbers of unsupervised exposures (e.g. Fig. 5E, right most points), suggesting that one or more of our assumptions and corresponding model components is not entirely accurate (see Discussion).

Given the surprising overall quantitative accuracy of the model predictions straight “out of the box”, we wondered if those predictions might somehow occur even for models that we had not carefully tuned to the initial (pre-exposure) human performance and the previously reported IT plasticity. That is, which components of the model are critical to this predictive success? We tested this in two ways (focusing here on experiment u1).

First, we changed the IT plasticity rate (α) to be either four times smaller or four times bigger than empirically observed in the prior monkey IT neurophysiology (solid gray lines) and re-ran the entire simulation procedure (above). In both cases, the predictions were now clearly different in magnitude than the observations (Fig. 5A). This result is arguably the strongest evidence that the single unit IT plasticity effects fully account for -- and do not over-account for -- the human unsupervised learning effects.

Second, we changed the two decoder hyperparameters (number of neural sites and number of training images) such that they were no longer correctly aligned with the initial human performance levels. Figure 5B illustrates the two-dimensional hyperparameter space, and the dashed line represents potential choices of the two hyperparameters that match human initial performance (the IT-to-COR-behavior matching manifold; Figure 3F). Regions above (or below) that manifold indicate hyperparameter choices where the decoders are better (or worse) performing than initial human performance. We found that the unsupervised learning effects predicted by the overall model (Fig. 5C, two black lines on top of each other corresponding to two choices of hyperparameters, black dots in Figure 5B) continued to well-approximate human learning effects. This was also true for other combinations of hyperparameters along the dashed black manifold in Fig. 5B (~10 combinations tested; results were similar to those shown in Fig. 5C,D,E, not shown). In other words, for model settings in which the overall model matched the initial human performance over all tasks, the predictions of the unsupervised learning effects remained similarly accurate.

In contrast, for choices of the two hyperparameters that did not match human initial performance, the unsupervised learning effect predicted by the overall model clearly differed from the observed human learning effect. Specifically, when the overall model starts off with “super-human” performance, it overpredicted the learning effect, and when it starts off as “sub-human” it underpredicted the learning effect.

In sum, it is not the case that any decoders will produce the correct predictions – proper setting of the IT-to-COR-behavior linkage model is critical. It is important to note that we did <u>not</u> tune these hyperparameters based on fitting the unsupervised effects in Fig. 2 – they were derived in accordance with prior work that did not involve any unsupervised learning (Majaj et al., 2015).

### The unsupervised learning effect depended on the initial task difficulty

So far, we have established a quantitative overall model that quite accurately predicted the direction, magnitude and time course of learning effects resulting from a range of unsupervised exposure manipulations. For each of those tests, we focused on object discrimination tasks that had an intermediate level of initial task difficulty (face discrimination tasks with initial d’ around 2.0), so that we had dynamic range to see both increases and decreases in performance (e.g. Fig. 2). However, we noticed that our IT plasticity rule seemed to imply that those learning effects would depend on the strength of the initial selectivity of individual IT neural sites for the exposed objects (i.e. the initial P vs. N response difference). The intuition is that this response difference is the driving force for IT plasticity updates (e.g. no difference leads to no update, large difference leads to large update). This in turn implied that the learning effect size should depend on the initial task performance (d’).

To test for this dependence, we focused on the unsupervised size tolerance “breaking” manipulation (as in u1, Fig. 2A, but with 800 unsupervised “swapped” exposures; see Methods), and tested new sets of human subjects using a wide range initial task difficulties, ranging from subordinate object discriminations (low d’) to basic level object discriminations (high d’). We focused on 13 size-specific object discrimination sub-tasks with either small-medium size swapping exposure or medium-big size swapping exposure. Each subject received only one exposure variant (see Methods). For each exposure variant, 20-40 new human subjects were tested, and we quantified the unsupervised learning effect (“breaking”) as the change (from initial) in performance (relative to control objects, as in Fig. 2A).

Figure 6B showed that unsupervised learning effect plotted against pre-exposure task difficulty for all 13 object discrimination tasks. This result not only confirms that this unsupervised learning effect is observed for a range of object discriminations (e.g. not just face objects), but it also showed a relationship between task difficulty (d’) and the magnitude of that learning effect. In particular, for initially easy tasks (d’ > ~2.5) and initially difficult tasks (d’ < ~0.5), we observed a smaller learning effect than tasks with intermediate initial performance.

We found that our overall model quite naturally -- for the reasons outlined above -- predicted the smaller learning effect for initially difficult tasks (the left side of Fig. 6B). Notably, the model did not naturally predict the lack of observed learning effects for the initially easy tasks (high d’ of Fig. 6B). However, that discrepancy could be simply explained if we assume there is a lapse rate (Prins, 2012) in human performance (that is, a fraction of trials for which the subject simply guesses, regardless of the quality of the sensory-driven information). The intuition here is that, human subjects make task-independent mistakes (“lapses”), and even a low rate of random lapses, puts a ceiling on the d’ value that can be experimentally measured (Fig, 6A). It is important to point out the attentional lapse rate has a much stronger influence on easy tasks than hard ones. In the context of our learning experiments, this would mean that the underlying neural representation for easy tasks might indeed be changing a great deal (at least, that is what our current model predicts), but those changes cannot be measured as changes in human performance in the face of a lapse-rate induced performance measurement ceiling (e.g. an “underlying” d’ of 5 changes to a d’ of 3.5, but we measure a d’ of ~3 in both cases and thus report a d’ change of ~0). Fig. 6B shows the predictions of the overall model (identical to the model used in Fig. 5), with an assumed lapse rate of 9% (9% random choices; 95.5% ceiling performance accuracy). This lapse rate value is supported by the performance accuracy of our basic level tasks (easy), which has a mean accuracy of 89% and maximum accuracy of 95%.

## Discussion

The goal of this study was to ask if previously reported temporal-contiguity driven unsupervised plasticity in IT neurons accounts for temporal-contiguity driven unsupervised learning effects in humans.

To do that, we built an overall computational model to predict human performance change resulting from plasticity in individual IT neural site firing rates under the paradigm of unsupervised temporal contiguity exposure (temporally contiguous pairs of images). The overall model had three core components: A) a generative model of a baseline adult IT neuronal population, B) an IT-population-to-COR-behavior linking model (adopted directly from (Majaj et al., 2015)), and C) an IT plasticity rule that aimed to capture and predict how pairs of temporally associated images lead to updates in the (future) image-driven responses of each individual IT neural site. Each of these three model components was guided by and constrained by prior work to set its parameters, and the combined overall computational model thus had zero free parameters.

To test the overall model, we asked the model to predict the human performance changes for three separate unsupervised learning experiments, and compared those predictions with the human performance changes (averaged over human subjects) that we measured in those three experiments. We found that the direction, magnitude, and time course of those mean unsupervised learning effects were all quite well predicted by the overall model (but not perfectly predicted). We also found that the model could naturally explain the dependence of the measured unsupervised learning on initial object discrimination difficulty (when we assumed a reasonable behavioral lapse rate).

In sum, this work successfully establishes a quantitative linking model between the plasticity in individual IT neurons and human behavioral changes (both improvements and disruptions) for temporal-contiguity driven unsupervised learning. More broadly, its success supports the overarching hypothesis that temporally contiguous unsupervised learning builds neural representations that underlie robust (i.e. tolerant) core object recognition, even in adults.

We were somewhat surprised that the overall model did such an accurate job of predicting the human learning effects essentially from predicted updates on the responses of IT neural sites. This was surprising because the overall model implicitly assumes that: monkey IT and human IT are approximately identical (Kriegeskorte et al., 2008; Rajalingham et al., 2018), the linkage of IT to core object recognition behavior is approximately identical in monkeys and humans (as previously suggested, (Majaj et al., 2015)), human unsupervised IT plasticity is the same as monkey IT plasticity (we are not aware of any previous tests of human IT plasticity under these conditions), that little or no behaviorally relevant plastic changes occur in the mechanistic linkage between IT and behavior during or after unsupervised visual experience (at least not at the time scales of these experiments: 1.5 to 2 hours). Of course, the results here do not prove any of the above assumptions, but they argue that it is most parsimonious to assume that all of the above are correct assumptions until further experiments show otherwise.

We noted a small discrepancy between the predictions of the model and the human learning data at the longest exposure durations that we tested (1600 exposure; ~1.5 hour); see Fig. 5A,C), where the model predicted slightly stronger behavioral changes than measured. One possibility is that learning over long periods of unsupervised exposure involves more complicated mechanisms in addition to our simplified unsupervised IT plasticity rule. For example, perhaps the plasticity rate slows down as the subject fatigues in the experiment. Or perhaps the plasticity mechanisms involve some type of renormalization of the responses of each IT neuron to retain some selectivity to different objects, as motivated by prior theoretical work on temporal contiguity learning (Sprekeler et al., 2007; Wiskott & Sejnowski, 2002). Similarly, plasticity along the ventral stream could involve homeostatic range adjustment which is fundamental to individual neurons (Turrigiano & Nelson, 2004), as motivated by studies of LTP and LTD plasticity in V1 neurons (e.g. BCM rules (Bienenstock, Cooper, & Munro, 1982; Toyoizumi, Pfister, Aihara, & Gerstner, 2005)). While we did not explicitly model any of these neural plasticity effects, they could be explored in future modeling studies.

## Tolerant object recognition and temporal-continuity driven unsupervised learning

Human (and monkey) visual object recognition is highly tolerant to object viewpoint, even under short, but natural, viewing durations of ~200 msec referred to as “core object recognition” (COR) (DiCarlo et al., 2012). Much evidence suggests that this ability derives from neural non-linear processing (and thus neural re-representation) of the incoming image along the ventral visual stream, and some ANN models have become reasonably accurate emulators of that non-linear processing and of its supported COR behavior (Cadieu et al., 2014; Krizhevsky et al., 2012; Kubilius et al., 2018; Yamins et al., 2014). However, because the “learning” of those models is highly non-biological (in the sense that millions of labeled images are used to explicitly supervise the learning), a key question remains completely open: how does the ventral stream develop its non-linear processing strategy?

One proposed idea is that, during post-natal development and continuing into adulthood, naturally-occurring temporally continuous visual experience can implicitly instruct plasticity mechanisms along the ventral stream that, working together, lead to the transform-invariant object representation (Berkes & Wiskott, 2005; Einhäuser, Hipp, Eggert, Körner, & König, 2005; Földiák, 1991; G. Wallis et al., 2009). Intuitively, the physics of time and space in our natural world constrains the visual experience we gain in everyday life. Because identity preserving retinal projections often occur closely in time, the spatiotemporal continuity of our viewing experience could thus be useful to instructing the non-linear processing that in turn supports highly view-tolerant object recognition behavior. Under this hypothesis, objects do not need to be labeled per se, they are simply the sources that statistically “travel together” over time.

We are not the first to propose this overarching hypothesis or variants of it, as the theoretical idea dates back to at least ~1960 (Attneave, 1954; Barlow, 1961). Földiák suggested that the internal representation should mimic physical entities in real life which are subject to continuous changes in time (Földiák, 1990, 1991). This process is purely unsupervised and achieves transformation invariance by extracting slow features from quickly varying sensory inputs (Berkes & Wiskott, 2005; Sprekeler et al., 2007; Wiskott & Sejnowski, 2002). A range of mathematical implementations of learning rules (Berkes & Wiskott, 2005; Földiák, 1991; Isik, Leibo, & Poggio, 2012; Körding, Kayser, Einhäuser, & König, 2004; Wiskott & Sejnowski, 2002) all include variants of this same conceptual idea: to achieve response stability of each neuron over time (while also maintaining response variance over the full population of neurons). Various synaptic plasticity mathematical rules and associated empirical observations support this form of unsupervised learning: Hebbian learning (D.O. Hebb, 1949; Földiák, 1991; Lowel & Singer, 1992; Paulsen & Sejnowski, 2000), anti-Hebbian Learning (Földiák, 1990; Mitchison, 1991; Pehlevan, Sengupta, & Chklovskii, 2017), BCM rule (Bienenstock et al., 1982; Toyoizumi et al., 2005) and spike-timing dependent plasticity (Caporale & Dan, 2008; Markram et al., 1997; Rao & Sejnowski, 2001). This prior work showed that unsupervised learning of neural representations of objects through temporal continuity was possible, at least in theory.

Human psychophysics studies have provided empirical evidence supporting the role of unsupervised temporal contiguity plasticity in visual object recognition. Wallis and Bulthoff found that unsupervised exposure to temporal image sequences of different views of different faces led performance deficits compared to sequences of the same face (Wallis & Bülthoff, 2001; Guy Wallis & Bülthoff, 1999). They also pointed out that these results were only observed in similar face pairs (i.e. low d’) rather than very distinct faces (i.e. higher d’). Cox et al showed that ‘swapped’ unsupervised experience of pairs of images across saccades could reduce (“break”) position tolerance of object discrimination (Cox et al., 2005). Balas and Sinha showed that observing object motion can increase both generalization to nearby views and selectivity to exposed views (Balas & Sinha, 2008). These behavioral observations revealed that unsupervised temporal contiguity is constantly contributing to the tolerance of object recognition behavior, even in adults, and thus it must be inducing some kind of underlying neural changes somewhere in the brain.

Our human psychophysical results reported here extend this prior work in three ways. First, we measured the learning effects over prolonged periods of time, which allowed us to test for accumulation and saturation. Second, we found that the behavioral learning effect is reversible (Fig. 2C). Third, we found that this unsupervised learning effect depended on initial task difficulty, which might explain why some studies report stronger effects than others. For example, Wallis and Bulthoff found that the learning effects on view tolerance were only observed in similar face pairs rather than very distinct faces (Wallis & Bülthoff, 2001), and those similar face pairs have initial d’ that happens to reside in the mid-range where we predict/observe the largest behavioral effects (Fig. 6B). Third, and most importantly, we designed our unsupervised visual statistical manipulations in the same way as previous monkey neurophysiology experiments, which allowed us to quantitatively compare our human behavioral results with prior monkey neuronal results.

Because IT is, among other ventral stream areas, thought to most directly underlie object discrimination behavior (DiCarlo et al., 2012; Ito et al., 1995; Rajalingham & DiCarlo, 2019) and IT plasticity has been found in many studies (Baker, Behrmann, & Olson, 2002; Logothetis, Pauls, & Poggio, 1995; Messinger, Squire, Zola, & Albright, 2001), reviewed by (Op de Beeck & Baker, 2010), it is natural to ask if temporally contiguous unsupervised experience also leads to plastic changes in IT neurons. Miyashita and colleagues showed that neurons in the temporal lobe shape their responses during learning of arbitrarily-paired images such that each neuron’s response becomes more similar to images that were presented nearby in time (Miyashita, 1988, 1993; Naya, Yoshida, & Miyashita, 2003; Sakai & Miyashita, 1991). Li and DiCarlo directly tested the role of unsupervised visual experience in IT neuronal tolerance by manipulating the identities and properties of objects presented consecutively in time (Li & DiCarlo, 2008, 2010). They found that, over ~1.5 hours of unsupervised exposure of ‘swapped’ temporal statistics, the size and position tolerance of IT neuronal responses were significantly modified, and that these changes were not reward or task dependent (Li & DiCarlo, 2012). Qualitatively similar unsupervised temporal continuity dependent neuronal plasticity has also been observed in rodents (Matteucci & Zoccolan, 2020).

While that prior experimental work seemed qualitatively consistent with the overarching theoretical idea, it did not demonstrate that the behavioral learning effects could be explained by the IT neural effects. The results of our study here show that those two effects are quantitatively consistent with each other – the behavioral effects are almost exactly accounted for by the IT neural effects.

### Future directions

One future direction is to extend our current overall model to other modalities, like view invariance or position invariance. This could be done by collecting further psychophysical data, adding proper tuning kernels to the current generative IT model and using the same IT plasticity rule and decoding model. A second future direction is to extend our current model to other objects beyond the 64 that have been tested in monkeys and human. This could be achieved through testing new IT population responses to new and old objects and then embedding the new objects in the multi-dimensional gaussian model of the IT population representation space based on neuronal population response similarity. Alternatively, we can use image computable deep artificial neural network models that quite accurately predict ventral stream neuronal population responses (Kubilius et al., 2018; Daniel L K Yamins et al., 2014) and use the “IT” layer to build a much larger representation space of objects. A third future direction is to develop new unsupervised learning algorithms that implement some of the core ideas of temporal contiguity learning, but are scaled to produce high performing visual systems that are competitive with state-of-the-art neural network systems trained by full supervision. Many computational efforts have touched on this direction (Agrawal, Carreira, & Malik, 2015; Bahroun & Soltoggio, 2017; Goroshin, Bruna, Tompson, Eigen, & LeCun, 2014; Higgins et al., 2016; Kheradpisheh, Ganjtabesh, & Masquelier, 2016; Lotter, Kreiman, & Cox, 2016; Srivastava, Mansimov, & Salakhutdinov, 2015; Wang & Gupta, 2015; Whitney, Chang, Kulkarni, & Tenenbaum, 2016), and some are just beginning to make predictions about the responses along the ventral stream (Zhuang, Yan, Nayebi, & Yamins, 2019; Zhuang, Zhai, & Yamins, 2019). A key next step will be to put those full scale models to experimental test at both the neurophysiological and behavioral levels.

## Acknowledgments

This work was supported in part by a grant from the National Institutes of Health to JJD (2-RO1-EY014970- 06) and by the Simons Foundation (SCGB [325500] to JJD).

## Author contributions

X.J. and J.J.D. designed the research; X.J. performed the experiments and the modeling; X.J., H.H, and J.J.D. analyzed data; X.J. and J.J.D. wrote the paper.

## Methods

### Datasets from prior work

To build a quantitative linking model that predicts unsupervised learning effects in humans from neuronal response in IT, we used three experimental datasets: 1. Human data: human psychophysics performance data collected with Amazon Mechanical Turk; 2. IT population data: simultaneous recordings of 168 sites with multi-electrode Utah array recordings implanted in monkey IT (from a previous study (Majaj et al., 2015)); and 3. IT single-site learning data: multi-unit activity recorded with single electrodes in monkey IT (from a previous study (Li & DiCarlo, 2010)).

#### IT population dataset

Multi-electrode array (Utah arrays) recordings from two awake macaque monkeys provided 168 multi-unit IT neural sites to 64 objects (5460 high-variation naturalistic images) for modeling use. Image presentation was 100 ms, and each image was repeated between 25 and 50 times. Spike counts were binned in the time window 70–170 ms post stimulus presentation and averaged across repetitions, to produce a 5,760 by 168 neural response pattern array. The 64 exemplar objects come from eight categories (animals, boats, cars, chairs, faces, fruits, planes and tables). Images were generated by placing a single exemplar object on a randomly-drawn natural scene background, at a wide range of positions, sizes and poses. Images were presented at 8 deg diameter at the center of gaze to awake fixating animals in a rapid serial visual presentation (RSVP) procedure (horizontal black bars indicate stimulus-presentation period). See (Majaj et al., 2015) for details.

#### IT plasticity dataset

Physiology data of unsupervised IT plasticity effects measured in multi-unit activities recorded at each single electrode in macaque monkey IT cortex were reanalyzed from Li & DiCarlo, 2010 (n=42 MUA sites). We refer to these as “neural sites.” The firing rate of sorted units were tested in response to preferred (P) and non-preferred (N) objects, each presented at a range of sizes, and were re-tested after different amounts of unsupervised exposure to evaluate the effect of that exposure on those response measures. See (Li & DiCarlo, 2010) for details.

### Image generation

We used the same 3D object models as previous published IT-behavior study (Majaj et al., 2015) and applied the same rendering mechanism (ray-tracing software) to each 3D object while parametrically varying its position, rotation and size, and projected on a randomly chosen unique natural background (out of a pool of 130 images) to generate new test image examples. All images were achromatic. The ground truth of each image was the identity of the generating 3D model, and this was used to evaluate performance accuracy. This naturalistic image generation allows us to gain full control of all the object-related meta-data in the images while preserving a relatively natural core object recognition experience.

For each object, we pre-define a “baseline view” (i.e. exact center of gaze, size of ~2 degree or 1/3 of the diameter of the image, and canonical pose; see Methods of (Majaj et al., 2015; Rajalingham et al., 2018)). Variations in size, position and rotation are transformations relative to baseline view of the object. Since our focus here was unsupervised learning of size-tolerant object selectivity, we intentionally introduced more images that only vary in size to measure size tolerance. Medium-sized objects were the ‘baseline’ size (~2 deg). Small-sized objects were 0.5x of baseline (~1 deg). Big-sized objects were 2x that of baseline (~4 deg). All test images for different sizes were generated with random naturalistic backgrounds. We thus created a set of 240 “size test” images per object (i.e. 80 images per object at each of the three test sizes). The final test images were each 512 x 512 pixels and were always presented to the subject at a total extent of ~7 degree of visual angle at the center of gaze (as in the prior neurophysiology studies above).

To neutralize possible size-specific attentional effects and possible size-specific adaptation effects, we presented these “size test” images intermixed with other “cover” images of the same objects. These cover images were generated using mild variation in all of the object view parameters. Specifically, we sampled randomly and uniformly from the following ranges: [- 1.2°, +1.2°] for object position in both azimuth (h) and elevation (v); [-2.4°, +2.4°] for rotation in all 3 axes; and [x0.7, x1.3] for size. These cover images were mixed randomly with the “size test” images (above) at a ratio of 1 cover image per “size test” image to generate a set of psychophysical test images for each subject (illustrated in supplemental Figure 1B). The behavioral results from the cover images were not part of the analyses.

### Human psychophysics and analysis

All human experiments were done in accordance with the MIT Committee on the Use of Humans as Experimental Subjects. A total of 505 (174 subjects in Figure 2 and 331 subjects in Figure 6) subjects completed our tasks published through Amazon’s Mechanical Turk, an online platform where subjects can participate in non-profit psychophysical experiments for payment based on the duration of the task. Aspects of core object recognition performance were measured based on the behavioral report following each test image presentation (Rajalingham et al., 2018). Previous work compared the results of core object recognition tasks measured in the laboratory setting with controlled viewing with results measured via Amazon MTurk, and found virtually identical results (Pearson correlation 0.94±0.01; from (Majaj et al., 2015)).

Each behavioral experiment contained two types of phases: a *Test Phase* in which specific aspects of object discrimination performance were measured (see below), and an *Exposure Phase* in which pairs of temporally contiguous images were experienced (See Fig. 1). The main experiment consisted of five Test Phases (200 trials each; 6-8 min) and four interleaved Exposure Phases (400 exposure events each; 12-20min) that together allowed us to measure exposure-induced changes in size-specific object discrimination over time (total experiment time ranged from 90 min to 120 min).

### Test phase

Our goal was to measure the discriminability of targeted (exposed) pairs of objects at targeted (exposed) sizes (and, as references, we also measured discriminability for control object pairs and for target objects at a non-exposed size). Conceptually, each such discrimination sub-task (size-specific object discrimination task) is a generated set of images from object A at a specific size that must be discriminated from a generated set of images of object B at a specific size, and mapped to the same object at a medium size (e.g, See Fig. 1B, choice images). For clarity, we note that, given this design, the only variation in each of these sub-task image test sets was the image background. These size-specific sub-tasks were randomly interleaved with cover trials to disguise this underlying fact from the subject (see Supp. Fig. 1B).

To measure performance on each sub-task, we used a two-way forced alternative classification (2AFC) design. Each 2AFC trial started with a central fixation point. Subjects were requested to fixate the black fixation point, because the test image was always presented briefly at that location and they might miss it otherwise. After 500 ms, the fixation dot disappeared and a test image appeared centered at dot location (center of the screen) for 100ms, followed by the presentation of two “choice” images presented on the left and right of the screen (Figure 1A). One of the choice images always matched the identity (or category) of the object that was used to generate the test image and was thus the correct choice, and its location was randomly assigned on each trial (50% on the right and 50% on the left). After mouse-clicking a choice image, the subject was given another fixation point (i.e. the next Test Phase trial began). No feedback on correctness of the choice was given.

To measure size-specific discrimination performance, we created size-specific 2AFC sub-tasks. Specifically, each sub-task was a balanced (i.e. 50% / 50%) set of size-specific test images generated from object A and object B (see above), and the two choices presented after each test image were “clean” examples of object A and object B at a standard (“medium”) size (see Fig. 1A).

For each subject, the test images were pseudorandomly drawn from the a test image pool that contained the desired number of ‘size test’ images and cover images (see Supp. Fig. 1B). Among the 200 trials (50 test image of each test object; 4 objects in total), 40% contained the “size test” images (20 for each object; 10 for small and 10 for big), 10% contained baseline views (medium size; 5 for each object), and the remaining 50% test images were “cover images” (see above) that were not used in analyses (see Supp. Fig. 1B for example test images). The number of test images for target and control object pairs were thus balanced. The number of test images for small and big sizes were also balanced regardless of exposure type. As a result, for each subject, we created six size-specific 2AFC sub-tasks in total (3 different sizes for each object pair) regardless of exposure type. The number of test images for target and control face pairs at different sizes in each test phase is specified in Supp. Fig. 1B.

To evaluate exposure induced learning effects, we only calculated the discrimination performance of three exposure-relevant sub-tasks (pre-planned, see Fig. 1B): 1. The sub-task with exposed (target) objects at the exposure manipulated size (Fig. 1b, red or blue d’); 2. the sub-task with non-exposed (control) objects at the exposure manipulated size (Fig. 1b, black d’) ; 3. the sub-task with exposed (target) objects at the non-manipulated size (Fig. 1b, dashed black d’). For example, one subject might have been randomly assigned to: exposure type = (experiment u1, swapped condition), target size = (big size), target objects = (face A, face B), control objects = (face C, face D). In this example, each Test Phase aimed to measure performance (d’) on three specific sub-tasks: [face A big vs. face B big], [face C big vs. face D big] and [face A small vs. face B small].

The sizes of the subject groups are provided in Results. The test trials for size and objects were always balanced in each subject group. In Figure 2, the subject groups differ in the exposure type (3 subject groups). In each of these three groups, the target exposure size was the big size, and within each group, the specific face objects for target and control were randomly selected for each subject. In Figure 6, the 13 subject groups correspond to the 13 subtasks that were targeted for exposure (see below). Within each subject group, the targeted type of object (i.e. face or basic level) and the targeted exposure size (i.e. medium-big or medium-small) was the same for all subjects, and within each group, the specific objects for target and control were randomly selected within the targeted type.

We computed the d’ for each exposure-relevant sub-task (typically three d’ values for each subject group; see Fig. 1B) based on the population (pooled) confusion matrix of the entire subject group. For each sub-task, we constructed a 2×2 confusion matrix by directly filling the behavioral choices into hit, miss, false alarm and correct rejection according to the stimuli and response of each trial (Fig. 1B). From the pooled confusion matrix, we computed the d’ for each sub-task. We used standard signal detection theory to compute d’s from the confusion matrix (d’ = Z(TPR) – Z(FPR), where Z is the inverse of the cumulative Gaussian distribution function, and TPR and FPR are true positive and false positive rates respectively). The d’ value was bounded within -7.4 to 7.4 (via an epsilon of 0.0001). The mean d’ for each sub-task of each subject group was determined by averaging the d’ calculated from each bootstrapped subjects sample (which is converges to the d’ of the pooled confusion matrix). The errorbar of performance (SEM) was estimated as the standard deviation of d’ over all bootstrap samples (1000 samples in each case),

### Exposure phase

Each exposure trial (a.k.a. exposure “event”) in the exposure phase was intended to deliver a pair of temporally-contiguous images at the center of gaze. Each trial initiated with the presentation of a small, central black dot (~0.5 deg), and the subject was required to mouse-click on that dot (this is intended to naturally bring the center of gaze to the dot). Immediately after a successful mouse-click (within 0.5 deg of the dot), two images were presented sequentially at the location of the black dot. Each image was shown for 100ms with no time lag between them. After the event, the black dot reappeared at a new, randomly-chosen location (out of 9 possible locations) on the screen (i.e. the next exposure trial began). The details of those images are described below in the context of the specific experiments carried out.

Because we here focused on the effects of unsupervised exposure events on size tolerance, the size of object in each of the two sequential images was always different and always included the medium (“baseline”) size: either big-sized objects paired with medium-sized objects, or small-sized objects paired with medium-size objects. In either variant, the order of those two images was counterbalanced, as in (Li & DiCarlo, 2010) (e.g.. approximately half of the events transitioned from medium to big objects and the other half from big to medium objects; signified by the double headed arrows in Fig. 1B).

### Flavors of unsupervised exposure

Following prior work (Cox et al., 2005; Li & DiCarlo, 2010; G Wallis & Bülthoff, 2001), there are two basic flavors of unsupervised exposure. The first flavor is referred to as the swapped exposure, in which the two images within each exposure event are generated from *different* objects (here, at different sizes). Based on prior work (Cox et al., 2005; G Wallis & Bülthoff, 2001) this exposure flavor is expected to gradually “break” (disrupt) size tolerance discrimination of those two objects. The second flavor is non-swapped exposure, in which the two images are generated from the *same* object (here, at different sizes). While this has been less studies in human psychophysics, based on prior IT neurophysiology results (Li & DiCarlo, 2010), this exposure flavor is expected to gradually build size tolerant discrimination of those two objects.

### Experimental designs

Our main experimental goal (Fig, 2 results) was to test the directions, magnitudes and temporal profiles of changes In size-tolerant object discrimination (assessed in the Test Phases, see above) resulting from different types of unsupervised exposure conditions (Figure 1&2). To do that, we deployed the two flavors (above) in three types of unsupervised experience types (u), and each subject was tested in only one of those three types. The first type (u1) was a series of swapped exposure epochs (intuitively, this aims for maximal “breaking”). The second type (u2) was a series of non-swapped exposure epochs (intuitively, this aims for maximal “building”). The third type (u3) was two swapped exposure epochs followed by two non-swapped exposure epochs (intuitively, this aims to test the reversibility of the unsupervised learning).

Each type of experiment lasted for about 90 minutes and each consisted of 9 phases in total: 5 test phases (200 test images each) and 4 exposure epochs (400 exposure events in each epoch; Figure 1A). This experiment was done with face objects only in a total of 174 subjects over all conditions (u1 = 102 subjects, u2 = 36 subjects, u3 = 37 subjects).

Our secondary experimental goal was to study how learning effect depends on the perceptual similarity of the exposed objects. To do this, we chose pairs of objects to cover a wide range of initial discrimination difficulties. Intuitively, it is easier to discriminate and elephant from a pear, than it is to discriminate an apple from a pear. Specifically, we chose a total of 13 size-specific object pairs selected from a set of 8 face objects (n=10 pairs) and 6 basic level objects (n=3 pairs). For subjects being exposed to faces, the control objects were also faces; for subjects exposed to basic level objects, the control objects were other basic level objects. These 13 pairs were selected based on pilot experiments that suggested that they would span a range of initial discrimination performance. Indeed, when tested in the full experiment (below), we found that mean human initial discrimination difficulties ranged broadly (d’ range: 0.4 to 2.6, based on the first Test Phase). We thus ran 13 groups of subjects (i.e. one group per target object/size pair) with ~20-40 subjects per group. Because the goal here was to test the *magnitude* of size-specific learning (not the time course), we tested only the “swapped” flavor of unsupervised experience using just one long exposure epoch (consisting of 800 exposure events). Each subject was exposed with only one pair of objects, and was exposed to one size variant of the exposure: small-medium size swapping or medium-big size swapping. We bracketed that unsupervised exposure with one pre-exposure Test phase (200 trials) and one post-exposure Test phase (also 200 trials). The learning effect was always measured at the exposed size (e.g. if exposed with small-medium swapping, the learning effect was measured as the performance change of small size discrimination task of the exposed object pair), subtracting the performance change for control object pair at the exposed size (all exactly analogous to Fig. 2A).

### Generative IT model

We modeled the IT neuronal population response based on the IT population dataset collected from monkey IT cortex with a Multi-Dimensional Gaussian (MDG) model. This model assumes that the distribution of IT population response (the distribution the mean responses of individual IT neurons to all images of an object) to each object is Gaussian-like. We tested this hypothesis with a normality test and found 81.25% (52 out of 64 distributions for 64 objects) of the IT population response distributions were Gaussian (reject when p<0.01). This MDG model preserves the covariance matrix of neuronal responses to all 64 objects that have been tested in monkey IT cortex. A random draw (of a

64×1 vector) from the MDG is conceptualized as the average response (over image repetitions) of a simulated IT recording site to each of the 64 objects. To generate the simulated IT tuning over changes in object size, we multiplied (outer product) that 64×1 vector with a randomly chosen size tuning kernel (1×3 vector) that was randomly sampled from a batch of size tuning curves that we had obtained by fitting curves to real IT responses across changes in presented object size (n=168 recording sites; data from (Majaj et al., 2015)). This gives rise to perfectly size tolerant simulated IT neurons (i.e. by construction, the tuning over object identity and over size are perfectly separable). To introduce more biological realism and to approximate the fact that each image is presented on a random background, we randomly jittered each value in the 64×3 matrix by a zero mean, iid shift of each matrix element (drawn from the distribution of variance across image exemplars for each object from IT neural data; σ^2^_clutter_). Given this procedure, we could generate a potentially infinite number of simulated IT neurons and their (mean) responses to each and every image condition of interest. We verified that, even with the simplifying assumptions imposed here, the population responses of simulated IT populations were quite similar to the actual IT neural population responses (in the sense of image distances in the IT population space, see Results).

To generate a hypothetical IT neural (model) population, we simply repeated the above process to obtain the requested number of model neurons in the simulated population (note: the MDG and the size tuning kernel pool was always fixed). In addition, when we “recorded” from these neurons (e.g. in Fig. 3A), we additionally added response “noise” that was independently drawn on each repetition of the same image (σ^2^_repeats_; mean zero, variance scaled with the mean to approximate known IT Poisson repetition “noise”).

### IT-to-behavior linking model

To generate behavioral performance predictions from model IT population responses, we applied a previously defined IT-to-recognition-behavior linking model (Majaj et al., 2015). In that study, the authors used actual IT neural population responses to show that a set of possible IT-to-behavioral linking models could each accurately describe and predict human performance on all tested recognition tasks within the reliability limits of the data. We here used one of the simplest, most biological plausible of those models --a linking model that seeks to infer the test image’s true label by computing the Pearson correlation between the mean IT population response to each possible object class (computed on the IT response to the training images) and the IT population response evoked by the current test image (note, test images are never used in the training of decoders). In other words, the model’s “choice” of object category for each test image was taken to be the choice object whose (simulated) IT population mean (over the training images) was closest to the population vector evoked by the current test image. The only difference from the prior work (Majaj et al., 2015) is that here we used simulated IT neurons (see Generative IT model, above) to drive the “behavior” of the model. (Note that the linking model has two key hyperparameters (see Results) and, for each simulation run, we held those constant.)

Since the model (IT population + linking model) could now be treated as a behaving “subject,” we analyzed the behavioral choices in exactly the same way as the actual human behavioral choices to arrive at d’ values that could be directly compared (i.e. generate a confusion matrix for each 2AFC sub-task, see above).

Similarly, to test a new model “subject,” we simply generated an entirely new IT model population (see above) and then found the parameters of the IT-to-behavior linking model for that subject.

To simulate human lapses (see Results), we introduced a (fixed) percentage of trials in each of the “behavioral” confusion matrices where model responses were randomly chosen. Note, when initial d’ is below ~2, the lapse rate most consistent with the data (9%) has little influence on measurable performance (see Fig. 6A) and thus only a minor effect on the model in Fig, 5.. Therefore, all predictions in Figure 5 were made with 0% lapse rate.

### Unsupervised IT plasticity model

We built a descriptive (non-mechanistic) learning rule with the same fundamental concept as previous computational models of temporal continuity learning (Földiák, 1990, 1991; Sprekeler et al., 2007; Wiskott & Sejnowski, 2002), except its mathematical implementation. In our setup, there are always only two images in each exposure event (a leading image and a lagging image). Our plasticity rule states that, after each exposure event, the modification of the mean firing rate response to the leading image is updated as follows:

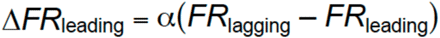

This plasticity rule tends to reduce the response difference between two exposed images (i.e. it tends to create response stability over time, assuming that the statistics of the future are similar). In our overall model, we apply this plasticity rule to each and every simulated IT neuron (true) after each and every exposure event. Note that, under repeated exposure events, the firing rate to all images will continue to change until there is no difference in responses to the leading and lagging images, which means the responses will eventually reach a steady state.

Compared with previous plasticity rules (e.g. Hebbian rule) for temporal continuity learning, our plasticity rule is relatively simple. Our plasticity rule updates each IT unit’s output firing rate directly rather than its input weights (Földiák, 1991). Based on immediate activity history, our learning rule continuously changes each unit’s output by pulling its responses to consecutive images closer until reaching steady state. This learning rule has several features. First, it is temporally asymmetric, which means the direction of rate change of the leading image depends on the sequence of leading and lagging image. In another word, the response to the lagging image is going to pull the response to the leading image towards it. However, since our experiments randomized the leading and lagging images on each exposure trial, this results is a change in the response to both images rather than an asymmetric change. Second, the effect of our plasticity rule is constrained to exposed image pairs and ignores any correlation in the neural representation space. Even though we do not yet have experimental data to accurately generalize the plasticity rule further than what has been presented in this paper, it is potentially generalizable to other types of tolerance (position, pose) and to other exposure paradigms.

The plasticity rate that best matches neural data is 0.0016 nru per exposure event (nru=normalized response units). The normalized response is calculated by Δ(P-N)/(P-N), where P and N represent the z-scored firing rate (across all objects) to preferred and non-preferred objects. Z-score is measured in terms of standard deviations from the mean. Therefore, 1 normalized response unit is 1 std of the response (firing rate) distribution across all tested objects. Since the mean multi-unit firing rate is 90±23 spk/sec (std across objects) for the IT population across 64 objects, we estimate that one nru is ~23 spk/sec. Therefore, 0.0016 nru corresponds to a firing rate change of ~0.035 spk/sec per exposure event, which means that ~30 exposure events of this kind would give rise to 1 spk/sec change in P vs. N selectivity.

**Supplementary figure 1.**
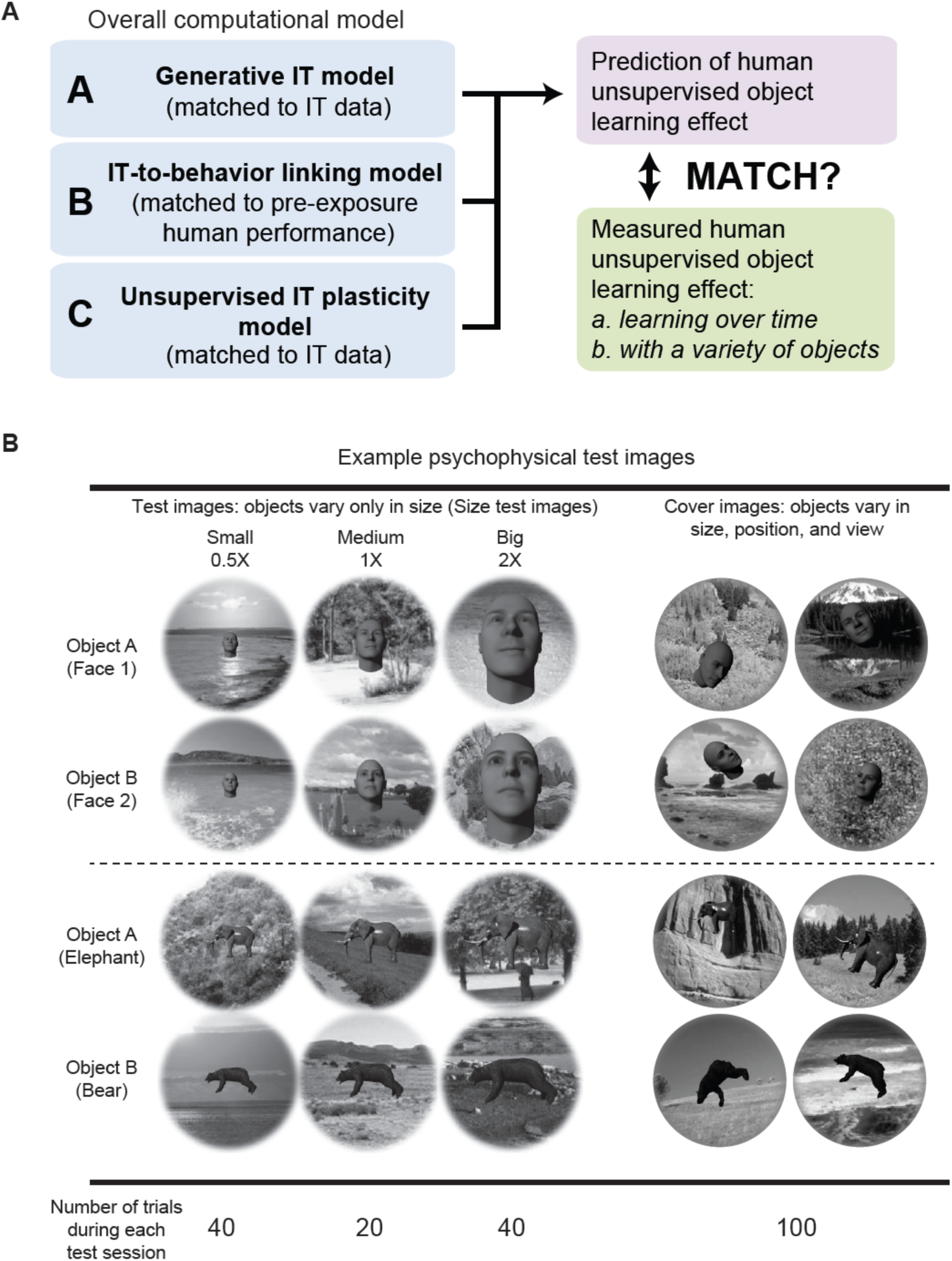
Paper outline and example test images. **A**) Outline diagram. **B**) Example test images and number of different types of test images per test phase (200 trials total).

## References

Afraz, A., Boyden, E. S., & DiCarlo, J. J. (2015). Optogenetic and pharmacological suppression of spatial clusters of face neurons reveal their causal role in face gender discrimination. Proceedings of the National Academy of Sciences of the United States of America, 112(21), 6730–6735. https://doi.org/10.1073/pnas.1423328112

Agrawal, P., Carreira, J., & Malik, J. (2015). Learning to See by Moving. Retrieved from http://arxiv.org/abs/1505.01596

Attneave, F. (1954). SOME INFORMATIONAL ASPECTS OF VISUAL PERCEPTION. Psychological Review (Vol. 61).

Bahroun, Y., & Soltoggio, A. (2017). Online Representation Learning with Single and Multilayer Hebbian Networks for Image Classification. Retrieved from http://arxiv.org/abs/1702.06456

Baker, C. I., Behrmann, M., & Olson, C. R. (2002). Impact of learning on representation of parts and wholes in monkey inferotemporal cortex. Nature Neuroscience, 5(11), 1210–1216. https://doi.org/10.1038/nn960

Balas, B., & Sinha, P. (2008). Observinga Object Motion Induces Increased Generalization and Sensitivity. Perception, 37(8), 1160–1174. https://doi.org/10.1068/p6000

Barlow, H. B. (1961). Possible Principles Underlying the Transformations of Sensory Messages.

Berkes, P., & Wiskott, L. (2005). Slow feature analysis yields a rich repertoire of complex cell properties. Journal of Vision, 5(6), 9–9. https://doi.org/10.1167/5.6.9

Bienenstock, E. L., Cooper, L. N., & Munro, P. W. (1982). Theory for the development of neuron selectivity: Orientation specificity and binocular interaction in visual cortex. Journal of Neuroscience, 2(1), 32–48. https://doi.org/10.1523/jneurosci.02-01-00032.1982

Cadieu, C. F., Hong, H., Yamins, D. L. K., Pinto, N., Ardila, D., Solomon, E. A., … DiCarlo, J. J. (2014). Deep Neural Networks Rival the Representation of Primate IT Cortex for Core Visual Object Recognition. PLoS Computational Biology, 10(12). https://doi.org/10.1371/journal.pcbi.1003963

Caporale, N., & Dan, Y. (2008). Spike Timing– Dependent Plasticity: A Hebbian Learning Rule. Annual Review of Neuroscience, 31(1), 25–46. https://doi.org/10.1146/annurev.neuro.31.060407.125639

Cox, D. D., Meier, P., Oertelt, N., & DiCarlo, J. J. (2005). “Breaking” position-invariant object recognition. Nature Neuroscience, 8(9), 1145–1147. https://doi.org/10.1038/nn1519

D.O. Hebb. (1949). The Organization of Behavior. Wiley: New York. https://doi.org/10.1016/s0361-9230(99)00182-3

DiCarlo, J. J., & Cox, D. D. (2007). Untangling invariant object recognition. Trends in Cognitive Sciences, 11(8), 333–341. https://doi.org/10.1016/j.tics.2007.06.010

DiCarlo, J. J., Zoccolan, D., & Rust, N. C. (2012). How Does the Brain Solve Visual Object Recognition? Neuron, 73(3), 415–434. https://doi.org/10.1016/j.neuron.2012.01.010

Einhäuser, W., Hipp, J., Eggert, J., Körner, E., & König, P. (2005). Learning viewpoint invariant object representations using a temporal coherence principle. Biological Cybernetics, 93(1), 79–90. https://doi.org/10.1007/s00422-005-0585-8

Földiák, P. (1990). Forming sparse representations by local anti-Hebbian learning. Biological Cybernetics, 64(2), 165–170. Retrieved from http://www.ncbi.nlm.nih.gov/pubmed/2291903

Földiák, P. (1991). Learning Invariance from Transformation Sequences. Neural Computation, 3(2), 194–200. https://doi.org/10.1162/neco.1991.3.2.194

Goroshin, R., Bruna, J., Tompson, J., Eigen, D., & LeCun, Y. (2014). Unsupervised Learning of Spatiotemporally Coherent Metrics. Retrieved from http://arxiv.org/abs/1412.6056

Hénaff, O. J., Goris, R. L. T., & Simoncelli, E. P. (2019). Perceptual straightening of natural videos. Nature Neuroscience. https://doi.org/10.1038/s41593-019-0377-4

Higgins, I., Matthey, L., Glorot, X., Pal, A., Uria, B., Blundell, C., … Lerchner, A. (2016). Early Visual Concept Learning with Unsupervised Deep Learning. Retrieved from http://arxiv.org/abs/1606.05579

Hung, C. P., Kreiman, G., Poggio, T., & DiCarlo, J. J. (2005). Fast Readout of Object Identity from Macaque Inferior Temporal Cortex. Science, 310(5749), 863–866. https://doi.org/10.1126/SCIENCE.1117593

Isik, L., Leibo, J. Z., & Poggio, T. (2012). Learning and disrupting invariance in visual recognition with a temporal association rule. Frontiers in Computational Neuroscience, 6, 37. https://doi.org/10.3389/fncom.2012.00037

Ito, M., Tamura, H., Fujita, I., & Tanaka, K. (1995). Size and position invariance of neuronal responses in monkey inferotemporal cortex. Journal of Neurophysiology, 73(1), 218–226. https://doi.org/10.1152/jn.1995.73.1.218

Kar, K., Kubilius, J., Schmidt, K., Issa, E. B., & DiCarlo, J. J. (2019). Evidence that recurrent circuits are critical to the ventral stream’s execution of core object recognition behavior. Nature Neuroscience, 22(6), 974–983. https://doi.org/10.1038/s41593-019-0392-5

Khaligh-Razavi, S.-M., & Kriegeskorte, N. (2014). Deep Supervised, but Not Unsupervised, Models May Explain IT Cortical Representation. PLoS Computational Biology, 10(11), e1003915. https://doi.org/10.1371/journal.pcbi.1003915

Kheradpisheh, S. R., Ganjtabesh, M., & Masquelier, T. (2016). Bio-inspired unsupervised learning of visual features leads to robust invariant object recognition. Neurocomputing, 205, 382–392. https://doi.org/10.1016/j.neucom.2016.04.029

Kingma, D. P., & Ba, J. (2014). Adam: A Method for Stochastic Optimization, 1–15. https://doi.org/http://doi.acm.org.ezproxy.lib.ucf.edu/10.1145/1830483.1830503

Körding, K. P., Kayser, C., Einhäuser, W., & König, P. (2004). How Are Complex Cell Properties Adapted to the Statistics of Natural Stimuli? Journal of Neurophysiology, 91(1), 206–212. https://doi.org/10.1152/jn.00149.2003

Kriegeskorte, N., Mur, M., Ruff, D. A., Kiani, R., Bodurka, J., Esteky, H., … Bandettini, P. A. (2008). Matching Categorical Object Representations in Inferior Temporal Cortex of Man and Monkey. Neuron, 60(6), 1126–1141. https://doi.org/10.1016/j.neuron.2008.10.043

Krizhevsky, A., Sutskever, I., & Hinton, G. (2012). ImageNet Classification with Deep Convolutional Neural Networks, 1–9.

Kubilius, J., Schrimpf, M., Nayebi, A., Bear, D., Yamins, D. L. K., & DiCarlo, J. J. (2018). CORnet: Modeling the Neural Mechanisms of Core Object Recognition. BioRxiv, 408385. https://doi.org/10.1101/408385

LeCun, Y., Boser, B., Denker, J. S., Henderson, D., Howard, R. E., Hubbard, W., & Jackel, L. D. (1989). Backpropagation Applied to Handwritten Zip Code Recognition. Neural Computation, 1(4), 541–551. https://doi.org/10.1162/neco.1989.1.4.541

LeCun, Yann, Bengio, Y., & Hinton, G. (2015). Deep learning. Nature, 521(7553), 436–444. https://doi.org/10.1038/nature14539

Li, N., Cox, D. D., Zoccolan, D., & DiCarlo, J. J. (2009). What Response Properties Do Individual Neurons Need to Underlie Position and Clutter “Invariant” Object Recognition? Journal of Neurophysiology, 102(1), 360–376. https://doi.org/10.1152/jn.90745.2008

Li, N., & DiCarlo, J. J. (2008). Unsupervised natural experience rapidly alters invariant object representation in visual cortex. Science (New York, N.Y.), 321(5895), 1502–1507. https://doi.org/10.1126/science.1160028

Li, N., & DiCarlo, J. J. (2010). Unsupervised natural visual experience rapidly reshapes size-invariant object representation in inferior temporal cortex. Neuron, 67(6), 1062–1075. https://doi.org/10.1016/j.neuron.2010.08.029

Li, N., & DiCarlo, J. J. (2012). Neuronal learning of invariant object representation in the ventral visual stream is not dependent on reward. Journal of Neuroscience, 32(19), 6611–6620. https://doi.org/10.1523/JNEUROSCI.3786-11.2012

Logothetis, N. K., Pauls, J., & Poggio, T. (1995). Shape representation in the inferior temporal cortex of monkeys. Current Biology : CB, 5(5), 552–563. Retrieved from http://www.ncbi.nlm.nih.gov/pubmed/7583105

Lotter, W., Kreiman, G., & Cox, D. (2016). Deep Predictive Coding Networks for Video Prediction and Unsupervised Learning. Retrieved from http://arxiv.org/abs/1605.08104

Lowel, S., & Singer, W. (1992). Selection of intrinsic horizontal connections in the visual cortex by correlated neuronal activity. Science, 255(5041), 209–212. https://doi.org/10.1126/science.1372754

Majaj, N. J., Hong, H., Solomon, E. A., & DiCarlo, J. J. (2015). Simple Learned Weighted Sums of Inferior Temporal Neuronal Firing Rates Accurately Predict Human Core Object Recognition Performance. Journal of Neuroscience, 35(39), 13402–13418. https://doi.org/10.1523/JNEUROSCI.5181-14.2015

Markram, H., Gerstner, W., & Sjöström, P. J. (2012). Spike-timing-dependent plasticity: a comprehensive overview. Frontiers in Synaptic Neuroscience, 4, 2. https://doi.org/10.3389/fnsyn.2012.00002

Markram, H., Lübke, J., Frotscher, M., & Sakmann, B. (1997). Regulation of synaptic efficacy by coincidence of postsynaptic APs and EPSPs. Science (New York, N.Y.), 275(5297), 213–215. Retrieved from http://www.ncbi.nlm.nih.gov/pubmed/8985014

Matteucci, G., & Zoccolan, D. (2020). Unsupervised experience with temporal continuity of the visual environment is causally involved in the development of V1 complex cells. Science Advances, 6(22), eaba3742. https://doi.org/10.1126/sciadv.aba3742

Messinger, A., Squire, L. R., Zola, S. M., & Albright, T. D. (2001). Neuronal representations of stimulus associations develop in the temporal lobe during learning. Proceedings of the National Academy of Sciences of the United States of America, 98(21), 12239–12244. https://doi.org/10.1073/pnas.211431098

Mitchison, G. (1991). Removing Time Variation with the Anti-Hebbian Differential Synapse. Neural Computation, 3(3), 312–320. https://doi.org/10.1162/neco.1991.3.3.312

Miyashita, Y. (1988). Neuronal correlate of visual associative long-term memory in the primate temporal cortex. Nature, 335(6193), 817–820. https://doi.org/10.1038/335817a0

Miyashita, Y. (1993). INFERIOR TEMPORAL CORTEX: Where Visual Perception Meets Memory. Annu. Rev. Neurosci (Vol. 16). Retrieved from www.annualreviews.org/aronline

Naya, Y., Yoshida, M., & Miyashita, Y. (2003). Forward processing of long-term associative memory in monkey inferotemporal cortex. The Journal of Neuroscience : The Official Journal of the Society for Neuroscience, 23(7), 2861–2871. https://doi.org/10.1523/JNEUROSCI.23-07 02861.2003

Oja, E. (1982). Simplified neuron model as a principal component analyzer. Journal of Mathematical Biology, 15(3), 267–273. https://doi.org/10.1007/BF00275687

Op de Beeck, H. P., & Baker, C. I. (2010, January). The neural basis of visual object learning. Trends in Cognitive Sciences. https://doi.org/10.1016/j.tics.2009.11.002

Paulsen, O., & Sejnowski, T. J. (2000). Natural patterns of activity and long-term synaptic plasticity. Current Opinion in Neurobiology, 10(2), 172–179. https://doi.org/10.1016/s0959-4388(00)00076-3

Pehlevan, C., Sengupta, A. M., & Chklovskii, D. B. (2017). Why do similarity matching objectives lead to Hebbian/anti-Hebbian networks? Retrieved from https://arxiv.org/pdf/1703.07914.pdf

Plaut, D. C., & Hinton, G. E. (1987). Learning sets of filters using back-propagation. Computer Speech and Language (Vol. 2). Retrieved from https://www.cs.toronto.edu/~hinton/absps/plautfilters.pdf

Prins, N. (2012). The psychometric function: The lapse rate revisited. Journal of Vision, 12(6), 25–25. https://doi.org/10.1167/12.6.25

Rajalingham, R., & DiCarlo, J. J. (2019). Reversible Inactivation of Different Millimeter-Scale Regions of Primate IT Results in Different Patterns of Core Object Recognition Deficits. https://doi.org/10.1016/j.neuron.2019.02.001

Rajalingham, R., Issa, E. B., Bashivan, P., Kar, K., Schmidt, K., & DiCarlo, J. J. (2018). Large-Scale, High-Resolution Comparison of the Core Visual Object Recognition Behavior of Humans, Monkeys, and State-of-the-Art Deep Artificial Neural Networks. The Journal of Neuroscience: The Official Journal of the Society for Neuroscience, 38(33), 7255–7269. https://doi.org/10.1523/JNEUROSCI.0388-18.2018

Rajalingham, R., Schmidt, K., & DiCarlo, J. J. (2015). Comparison of Object Recognition Behavior in Human and Monkey. The Journal of Neuroscience : The Official Journal of the Society for Neuroscience, 35(35), 12127–12136. https://doi.org/10.1523/JNEUROSCI.0573-15.2015

Rao, R. P. N., & Sejnowski, T. J. (2001). Spike-Timing-Dependent Hebbian Plasticity as Temporal Difference Learning. Neural Computation, 13(10), 2221–2237. https://doi.org/10.1162/089976601750541787

Riesenhuber, M., & Poggio, T. (1999). Hierarchical models of object recognition in cortex. Nature Neuroscience, 2(11), 1019–1025. https://doi.org/10.1038/14819

Rolls, E. T., & Stringer, S. M. (2006). Invariant visual object recognition: A model, with lighting invariance. Journal of Physiology-Paris, 100(1–3), 43–62. https://doi.org/10.1016/j.jphysparis.2006.09.004

Rust, N. C., & DiCarlo, J. J. (2010). Selectivity and Tolerance (“Invariance”) Both Increase as Visual Information Propagates from Cortical Area V4 to IT. Journal of Neuroscience, 30(39), 12978– 12995. https://doi.org/10.1523/JNEUROSCI.0179-10.2010

Sakai, K., & Miyashita, Y. (1991). Neural organization for the long-term memory of paired associates. Nature, 354(6349), 152–155. https://doi.org/10.1038/354152a0

Sprekeler, H., Michaelis, C., & Wiskott, L. (2007). Slowness: An Objective for Spike-Timing– Dependent Plasticity? PLoS Computational Biology, 3(6), e112. https://doi.org/10.1371/journal.pcbi.0030112

Srivastava, N., Mansimov, E., & Salakhutdinov, R. (2015). Unsupervised Learning of Video Representations using LSTMs. Retrieved from http://arxiv.org/abs/1502.04681

Toyoizumi, T., Pfister, J. P., Aihara, K., & Gerstner, W. (2005). Generalized Bienenstock-Cooper-Munro rule for spiking neurons that maximizes information transmission. Proceedings of the National Academy of Sciences of the United States of America, 102(14), 5239–5244. https://doi.org/10.1073/pnas.0500495102

Turrigiano, G. G., & Nelson, S. B. (2004). Homeostatic plasticity in the developing nervous system. Nature Reviews Neuroscience. European Association for Cardio-Thoracic Surgery. https://doi.org/10.1038/nrn1327

Wallis, G., Backus, B. T., Langer, M., Huebner, G., & Bulthoff, H. (2009). Learning illumination-and orientation-invariant representations of objects through temporal association. Journal of Vision, 9(7), 6–6. https://doi.org/10.1167/9.7.6

Wallis, G, & Bülthoff, H. H. (2001). Effects of temporal association on recognition memory. Proceedings of the National Academy of Sciences of the United States of America, 98(8), 4800–4804. https://doi.org/10.1073/pnas.071028598

Wallis, Guy, & Bülthoff, H. (1999). Learning to recognize objects. Trends in Cognitive Sciences, 3(1), 22– 31. https://doi.org/10.1016/S1364-6613(98)01261-3

Wang, X., & Gupta, A. (2015). Unsupervised Learning of Visual Representations using Videos. Retrieved from http://arxiv.org/abs/1505.00687

Whitney, W. F., Chang, M., Kulkarni, T., & Tenenbaum, J. B. (2016). Understanding Visual Concepts with Continuation Learning. Retrieved from http://arxiv.org/abs/1602.06822

Wiskott, L., & Sejnowski, T. J. (2002). Slow Feature Analysis: Unsupervised Learning of Invariances. Neural Computation, 14(4), 715–770. https://doi.org/10.1162/089976602317318938

Yamins, D. L. K., Hong, H., Cadieu, C. F., Solomon, E. A., Seibert, D., & DiCarlo, J. J. (2014). Performance-optimized hierarchical models predict neural responses in higher visual cortex. Proceedings of the National Academy of Sciences, 111(23), 8619–8624. https://doi.org/10.1073/pnas.1403112111

Yamins, Daniel L K, Hong, H., Cadieu, C. F., Solomon, E. A., Seibert, D., & DiCarlo, J. J. (2014). Performance-optimized hierarchical models predict neural responses in higher visual cortex. Proceedings of the National Academy of Sciences of the United States of America, 111(23), 8619– 8624. https://doi.org/10.1073/pnas.1403112111

Zhuang, C., Yan, S., Nayebi, A., & Yamins, D. (2019). Self-supervised Neural Network Models of Higher Visual Cortex Development. https://doi.org/https://doi.org/10.32470/CCN.2019.1393-0

Zhuang, C., Zhai, A. L., & Yamins, D. (2019). Local Aggregation for Unsupervised Learning of Visual Embeddings. Retrieved from http://arxiv.org/abs/1903.12355

